# Sensitive Detection of Structural Differences using a Statistical Framework for Comparative Crystallography

**DOI:** 10.1101/2024.07.22.604476

**Authors:** Doeke R. Hekstra, Harrison K. Wang, Margaret A. Klureza, Jack B. Greisman, Kevin M. Dalton

## Abstract

Chemical and conformational changes underlie the functional cycles of proteins. Comparative crystallography can reveal these changes over time, over ligands, and over chemical and physical perturbations in atomic detail. A key difficulty, however, is that the resulting observations must be placed on the same scale by correcting for experimental factors. We recently introduced a Bayesian framework for correcting (scaling) X-ray diffraction data by combining deep learning with statistical priors informed by crystallographic theory. To scale comparative crystallography data, we here combine this framework with a multivariate statistical theory of comparative crystallography. By doing so, we find strong improvements in the detection of protein dynamics, element-specific anomalous signal, and the binding of drug fragments.

## Introduction

Proteins and their assemblies mediate chemical catalysis, molecular transport, signal transduction, and the allosteric control of these processes. X-ray crystallography is rapidly expanding experimental access to these processes, driven by the advent of new X-ray sources including fourth-generation synchrotrons^1^ and X-ray free-electron lasers (XFELs)^2,3^, by new time-resolved methods such as mix-and-inject studies of enzyme catalysis^4-8^, and by high-throughput methods permitting screening of the interactions of thousands of molecules with proteins of interest^9-14^. These methods enable tracking of enzymatic intermediates in atomic detail, mapping of the concerted motions of proteins^10,15-17^, and the identification of protein-binding drug fragments (small-molecule templates for drug design). Each of these experiments is comparative and seeks to detect subtle changes in average structure. The success of such experiments hinges not only on billion-dollar instruments, but also on the sensitivity of the algorithms used to detect changes in structure from the X-ray diffraction data.

X-ray diffraction images obtained from macromolecular crystals consist of patterns of many small spots, or reflections. The true intensities of these reflections are proportional to the square amplitudes of the “structure factors”—the Fourier components of the electron density in the crystal. The observed intensities, however, also depend on a range of multiplicative factors (“scales”) due to the intensity and polarization of the incident beam, the volume of crystal exposed to the beam, intrinsic and beam-induced crystal defects, and absorption of the beam by surrounding material, air, and the detector itself. Neither the true reflection intensities nor these scales are directly observed, yet it is critical that before inferring changes in structure from changes in observed intensities, these intensities must be corrected. The correction procedure is known as scaling.

Approaches to scaling rely on comparison of redundant observation of equivalent reflections which should yield the same structure factor amplitude up to variation in the scale factor. When sample or beamtime are limited, the need for scaling to compare different conditions generates a tradeoff: one can either accurately estimate scales by maximizing redundancy for a few conditions, or observe many conditions but be left with residual systematic differences in the scales of each dataset. These remaining scaling inaccuracies are often addressed after the fact, for example by applying local scaling in SOLVE^18^, SCALEIT^19^, or the “Isomorphous Difference Map” utility of PHENIX (e.g., in ^20,21^).

Here we introduce a framework that enables statistically efficient comparison of related datasets. We first observe that the structures of proteins at different timepoints or under different chemical or physical conditions are often very similar, and therefore could provide nearly redundant measurements. To exploit this similarity, we adapted an algorithm we recently introduced for scaling and merging of X-ray diffraction data^22^. This algorithm, implemented in the software package *Careless*, applies to diffraction data from all predominant experimental approaches. *Careless* is based on approximate Bayesian inference combined with a neural network that learns scale factors from experimental metadata such as the position of each diffraction spot on the detector, Miller indices, and frame or crystal number. To adapt this approach to comparative applications, we developed a multivariate prior distribution that accurately captures correlations between related crystallographic datasets, and extend the *Careless* formalism to use such prior information. We show that this approach leads to strong improvements in the detection of signals in several examples—detection of protein dynamics from a polychromatic, time-resolved experiment, detection of specific elements in enzymes from serial XFEL and polychromatic synchrotron data, and the detection of bound drug fragments from a crystallographic screen. In forthcoming work^23,24^, we further show that this approach makes it possible to follow enzyme catalysis in a serial, rapid-mixing experiment.

### A statistical framework for comparative crystallography

Our statistical model is based on three concepts and explained in mathematical detail in the **Supplementary Information**: **First**, each structure factor, or Fourier component of the electron density, can be thought of as the sum of contributions from all the constituent atoms^25^. The sum of these contributions is well-approximated by the central limit theorem. The structure factors, which have real and imaginary components, therefore follow a bivariate normal distribution in the complex plane centered at the origin (**Figure 1a**, black arrows). The structure factor amplitudes, which correspond to the distance from the origin in **Figure 1a** (dashed circle), follow the well-known Wilson distribution^26^. **Second**, when comparing two related datasets, the contributions of corresponding atoms in the two crystals will be similar and therefore the sums of their contributions to their respective structure factors will be similar. When considered together, they will closely follow a multivariate normal distribution characterized by a correlation parameter, henceforth the *double-Wilson r*^27,28^, that quantifies the correlation between the real components of pairs of structure factors. Importantly, the joint distribution of complex structure factors ***F***_1_, ***F***_2_ can be factorized such that *P*(***F***_1_, ***F***_2_) = *P*(***F***_1_|***F***_2_) · *P*(***F***_2_) with *P*(***F***_1_|***F***_2_) only dependent on the difference in phases of ***F***_1_ and ***F***_2_. Integrating this unobserved phase difference yields a conditional probability density *P*(*F*_1_| *F*_2_) that follows the Rice distribution^28-30^ (where *F* = | ***F***|). We will refer to this statistical model as the double (or bivariate) Wilson model, and to the resulting joint distribution of structure factor amplitudes as the bivariate Wilson distribution.

**Figure 1:**
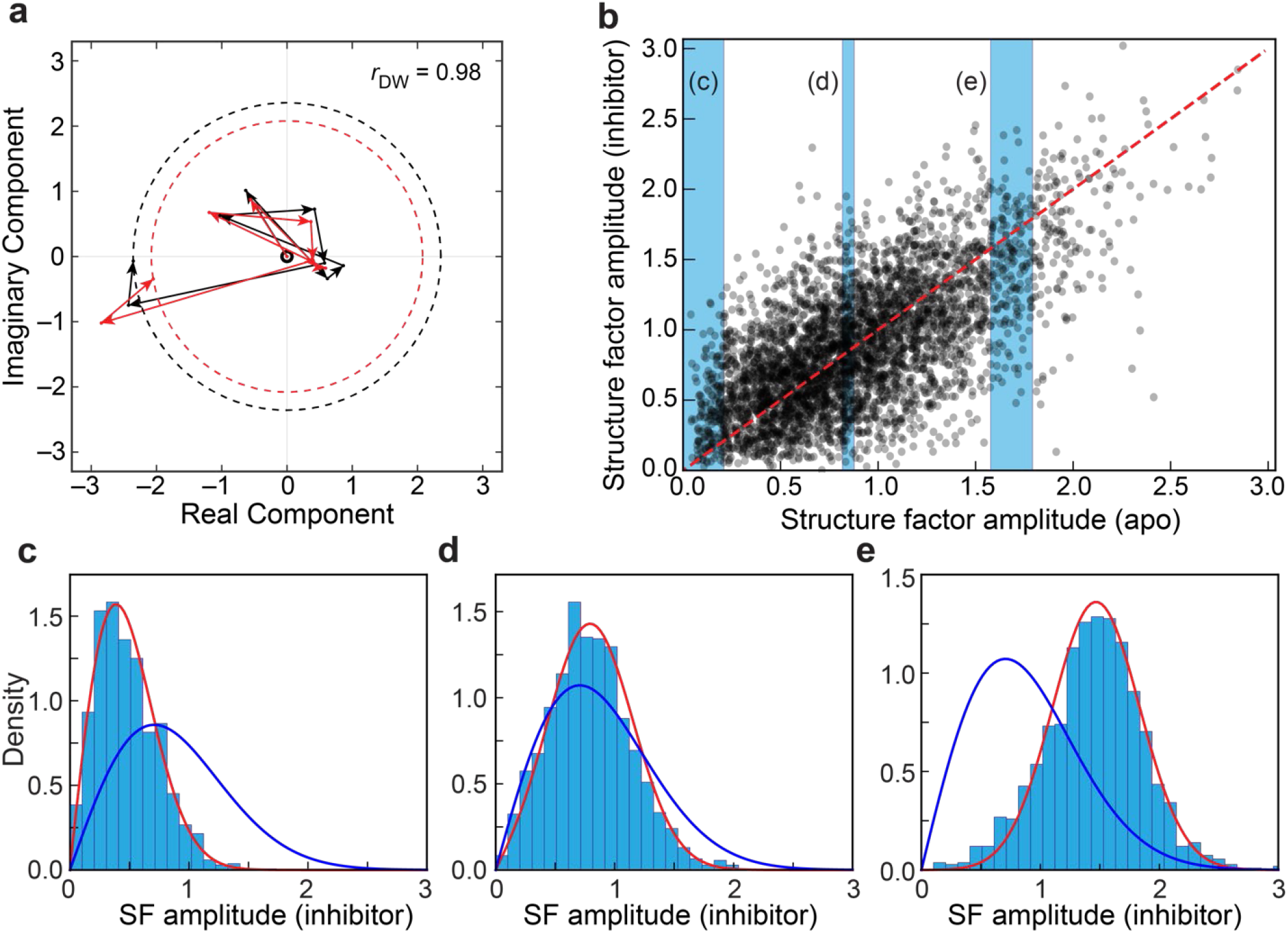
The double-Wilson model. **a)** In the double-Wilson model, the structure factors of related structures (black and red) are built up from pairs of random contributions (arrows) with correlation *r*_*D* w_ between their real components and likewise between their imaginary components. The experimentally observed amplitudes of the structure factors are described by the radii of the dashed circles. **b)** Scatter plot for structure factor amplitudes for isomorphous PTP-1B datasets for *apo* and inhibitor-bound forms (for a random subset of Miller indices). Blue slices indicate data points for which histograms are shown in panels c-e. **c-e)** Histograms for slices through panel b are better approximated by the Rice distributions (red) parametrized by a global *r*_*Dw*_ (here 0.85) than by the unconditional, univariate Wilson distribution (blue).

**Figure 1b** compares structure factor amplitudes for crystals of human Protein Tyrosine Phosphatase 1B (PTP-1B) in the presence and absence (apo) of the inhibitor TCS-401^31^. A fair degree of correlation is evident. A fit to the double-Wilson model (**Supplementary Notebook 3**) shows that *r* ≈ 0.85, with the implied Pearson correlation coefficient between structure factor amplitudes ≈ *r*^2^ (**Supplementary Notebook 2a**)^32^. Indeed, histograms of structure factor amplitudes for the inhibitor-bound data (**Figure 1c-e and Figure S1**) given the binned amplitudes of the apo data (blue shading in **Figure 1b**) are better fit by this model (red curves in **Figure 1c-e**) than by assuming statistical independence (blue curves). We likewise find that the bivariate Wilson distribution describes other comparative crystallography datasets well, including time-resolved X-ray crystallography experiments (**Figure S2**), replicate data sets across labs (**Figure S3**), and across temperatures (**Figure S4**), providing confidence that the formalism generally applies to related datasets.

**Third**, this approach does not generalize to more than two datasets, as it is not known how to integrate over the unknown phase differences. However, for many crystallographic experiments, it is reasonable to assume some conditional independence. For instance, when structure factor amplitudes A and C are both correlated with structure factor amplitudes B, their joint probability is often well described as the product *P*(*A, B, C*) ≈ *P*(*A*|*B*) · *P*(*C*|*B*) · *P*(*B*) rather than the exact expression *P*(*A, B, C*) = *P*(*A*|*B, C*) · *P*(*C*|*B*) · *P*(*B*), as long as *C* does not provide additional information about *A* not contained in *B*. Assumptions about conditional (in)dependence can be summarized in a Bayesian network or graphical model^33^ (**Figure 2a**). In this representation, each node represents a dataset, and each edge (arrow) represents conditional dependence. Whenever the resulting graph is acyclic, the resulting joint probability of structure factor amplitudes can be calculated analytically (**Supplementary Information**). We will refer to this distribution as the multivariate Wilson distribution. We note that conditional dependence is reciprocal, and the orientation of the displayed arrows is irrelevant for our purposes. Indeed, the expected correlations in many comparative crystallography experiments can be approximated by such networks (**Figure 2b**). For example, for a crystallographic time series, when estimating structure factors at time 3, it is a good assumption that we would not gain anything from knowing the structure factor amplitudes at time 1 if we already have access to their values at time 2. We further note that not all nodes need to have been observed—conditional independence mediated by a ‘dummy’ has found application in the analysis of multiple isomorphous replacement^34^.

**Figure 2.**
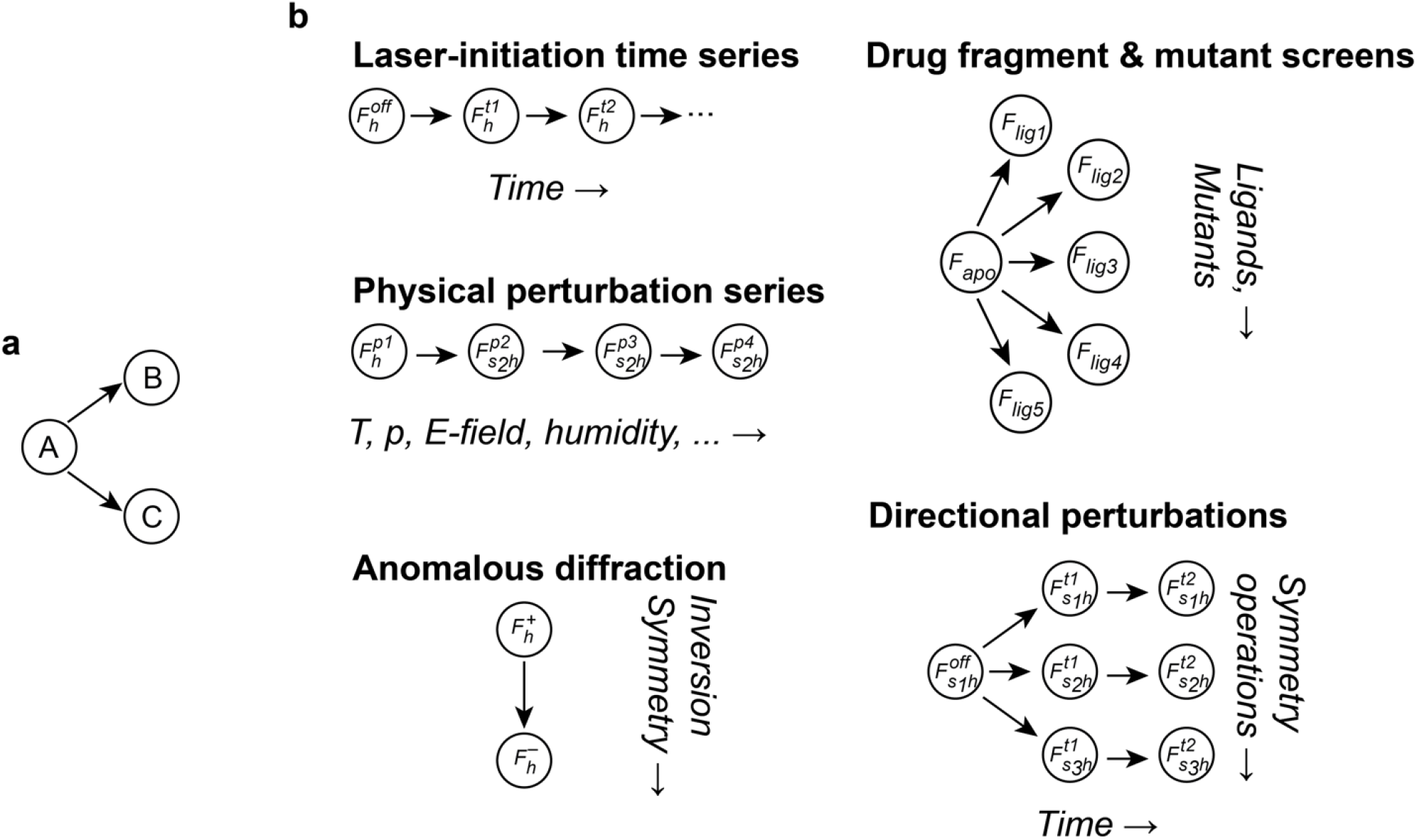
Modeling correlations of structure factors in comparative crystallography experiments. **a**. A graphical model (or Bayesian network) illustrating the relationship between three datasets (or, more generally, sets of structure factor amplitudes). Arrows indicate conditional dependence; the lack of an arrow between datasets B and C indicates conditional independence. **b**. Correlations in many experiments are well approximated by acyclic graphical models for which the joint prior distribution of structure factor amplitudes *F*_*h*_ can be calculated analytically. *t*_1_, *t*_2_, … represent timepoints; *s*_1_, *s*_2_, … represent symmetry operations; *p*_1_, *p*_2_, … represent perturbations due to, e.g., temperature *T*, pressure *p*, or electric field; *F*^+^ and *F*^−^ are reflections related by Friedel’s law.

### Scaling of related datasets

We recently introduced a Bayesian statistical framework for simultaneous inference of scales and structure factor amplitudes from reflection metadata (such as Miller indices, image number, position on the detector, and X-ray wavelength) and observed intensities^22^. Until now, estimation of each structure factor amplitude has been independently constrained by a Wilson prior distribution on the amplitudes. For comparative applications, this approach is inefficient: during scaling, more accurate scales enable more accurate estimates of structure factor amplitudes, and vice versa. It would be natural to incorporate a joint prior distribution on structure factor amplitudes during scaling that reflects the expected correlations between conditions. Different scenarios for a pair of related structure factor amplitudes *F*_1_, *F*_2_ illustrate the anticipated benefits of a multivariate approach: (i) if only noisy observations are available for *F*_1_ and *F*_2_, the prior should force estimates of *F*_1_ and *F*_2_ to be highly similar and, therefore, their difference small; (ii) when *F*_1_ can be estimated accurately, the anticipated similarity of the structure factor amplitudes will facilitate more accurate estimation of the scales for the other dataset by narrowing down the range of probable values of *F*_2_; (iii) when accurate observed intensities are available for both datasets, the anticipated similarity helps calibrate the relative scales in both datasets. With these advantages in mind, we implemented the multivariate Wilson distribution as a prior in Careless and show in four examples that structure factor differences can, indeed, be much better estimated upon scaling amplitudes with a multivariate prior. Operationally, our implementation expects users to supply the topology of conditional dependencies between datasets (**Figure 2b**) and an estimate of the double-Wilson parameter *r*.

### Laue anomalous diffraction

Although many X-ray sources are intrinsically polychromatic, most of their photons are typically discarded by use of monochromators. For studies of dynamics, however, it can be essential to use much brighter polychromatic pulses, giving rise to Laue diffraction. To assess whether the use of multivariate priors can improve the processing of Laue data, we first addressed the extraction of anomalous signal, which can be recovered from small differences between Bijvoet pairs—pairs of structure factors that are identical in the absence of element-specific electronic resonances (that is, anomalous effects) but that differ when such effects are present.

In this example, all data were collected on a single crystal of hen egg white lysozyme soaked with sodium iodide using polychromatic X-ray pulses (about 5% spectral bandwidth, from 1.02 to 1.20 Å wavelength; see **Methods**). As these data were obtained from a single crystal, one might expect little inconsistency in artifacts across the data and minimal benefit from imposing a bivariate prior on Bijvoet pairs. Nevertheless, we compared scaling and merging in Careless with a univariate Bayesian prior (i.e., the standard Wilson distribution) against a bivariate prior that assumes varying levels of correlation, *r*, between the Bijvoet mates. For consistency, we phased all sets of structure factor amplitudes using a reference model refined against monochromatic data collected on the same day (see **Methods**). In the resulting anomalous difference map, we find strong gains in peak heights of both iodide ions and sulfur atoms when using a bivariate prior (**Figure 3a, Figure S5a**), even though the anomalous scattering strength of sulfur is only about 1/4^th^ the scattering from a single electron (*f*′′ = 0.26 at the dominant wavelength of the incident X-ray spectrum, 1.04 Å). Importantly, the signal strength is maximal at an intermediate *r* of 0.999, with an increase in anomalous peak height of about 50% over the use of a univariate prior (**Figure 3b, Figure S5b**). Cross-validation measures of data processing quality (*CC*_1/2_ ^35^, *CC*_pred_ ^36^, *CC*_anom_^37^) are also improved by the introduction of the bivariate prior (**Figure 3c-d**).

**Figure 3.**
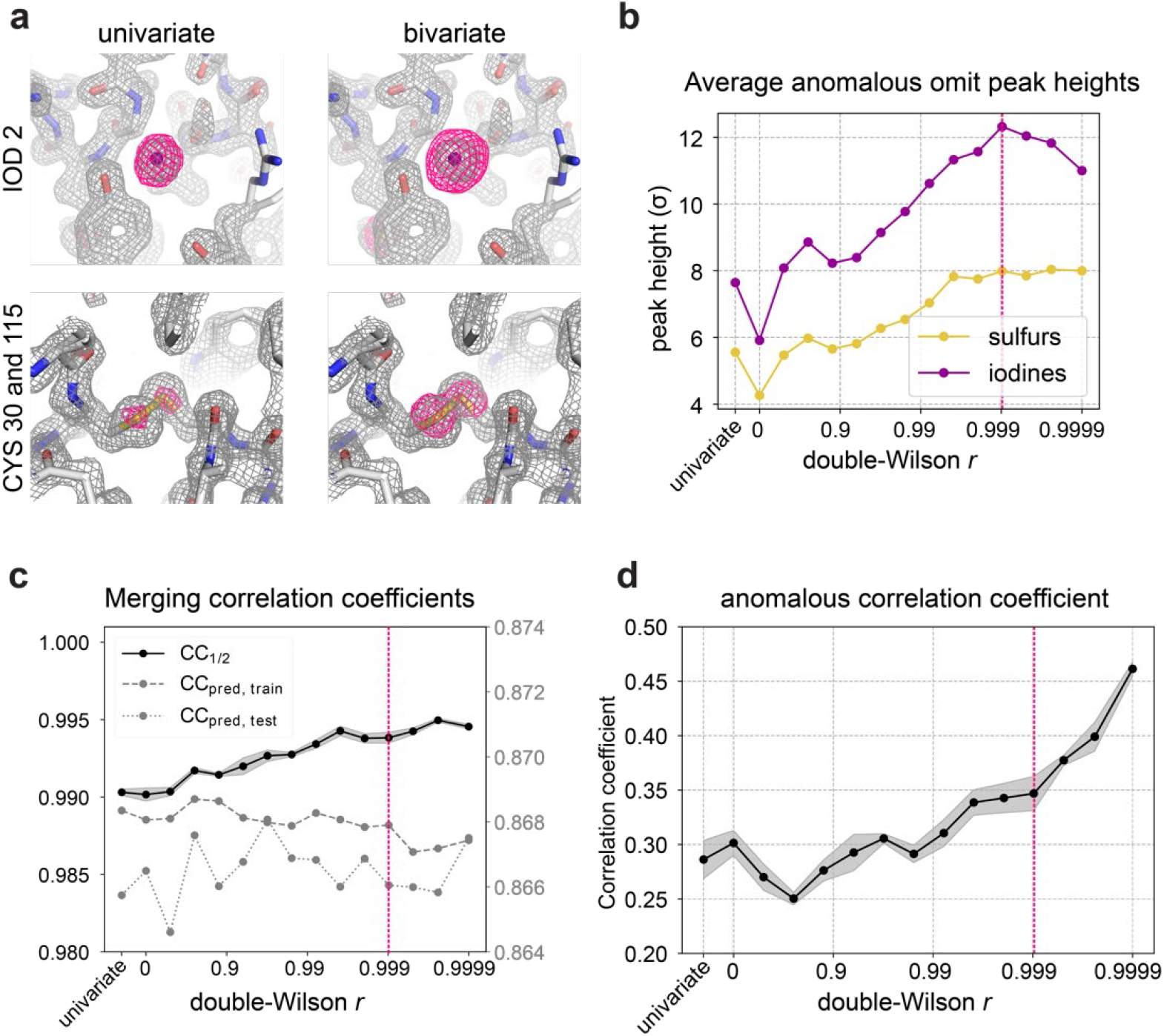
Use of a bivariate prior improves anomalous signal in a Laue diffraction experiment. **a)** Comparisons between anomalous omit peaks after scaling and merging with univariate or bivariate priors (*r*= 0.999 for the bivariate prior, marked in panels **b-d** with a dashed magenta line). **b)** Dependence of average iodine and sulfur anomalous peak height on *r*. **c)** Merging correlation coefficients of the lysozyme dataset across double-Wilson *r* values. The y-axis labels for the *CC*_1/2_ are on the left, and the y-axis labels for the *CC*_pred_ are on the right. A test set of 10% of observations were held out during scaling and merging to evaluate performance of the scaling model, yielding *CC*_pred, test_ for the test set, and *CC*_pred, train_ for the 90% of data used during scaling. The shaded confidence interval of the *CC*_1/2_ curve represents the standard deviation over three half-dataset repeats. **d)** Anomalous correlation coefficient of the lysozyme dataset across *r*. The shaded confidence interval represents the standard deviation over three half-dataset repeats.

### Time-resolved Laue diffraction

Laue diffraction at synchrotron sources provides access to the dynamics of proteins on timescales of 100 picoseconds and longer. To determine whether a bivariate prior can improve scaling of time-resolved signal, we processed a time-resolved Laue dataset collected on a single crystal of photoactive yellow protein (PYP) at BioCARS 14-ID at the Advanced Photon Source.

The light-induced *trans*-to-*cis* isomerization of the active site chromophore (**Figure 4a**) has been studied extensively^38-40^. Our dataset contains 20 images of PYP without laser exposure and 20 images 2 milliseconds after exposure to a blue laser pulse, at which point PYP is expected to adopt the pB1 state^40^. Using Careless, we jointly scaled the OFF and 2ms datasets while imposing a joint prior with varying levels of the correlation parameter, *r*. Merging statistics improve when scaling with a bivariate prior (**Figure 4b**). One such statistic is the correlation coefficient, *CC*_pred_, between observed intensities and those implied by the scaling model and the estimated structure factor amplitudes^22^. The gap between the *CC*_pred_ for training data and a cross-validation test set is reduced with increasing *r*. This means that the bivariate prior protects the scaling model against overfitting. The extensive literature on PYP enables us to quantify the quality of the resulting electron density difference maps: based on models of the ground-state and excited-state conformations from previous studies, we can calculate real-space correlation coefficients with a predicted difference map (that is, the correlation with an 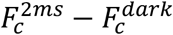 map based on conformations from Protein Data Bank (PDB) entry 1TS0, see ref. ^40^). At the best value of *r* (about 0.999), the real-space CC (Correlation Coefficient) of the observed difference map is more than 0.50 within 10 Å of the chromophore, about twice as strong as for an observed difference map obtained by scaling with a univariate prior (**Figure 4c**). Consistent with this quantitative assessment, the difference map near the chromophore is more readily visually interpretable after scaling with a bivariate prior (**Figure 4d**; see also **Figure S6**). We also attempted to place ON and OFF datasets on the same scale using local scaling or SCALEIT^41^ (**Methods**). In our hands, neither approach was effective (**Figure 4c**, on the left).

**Figure 4:**
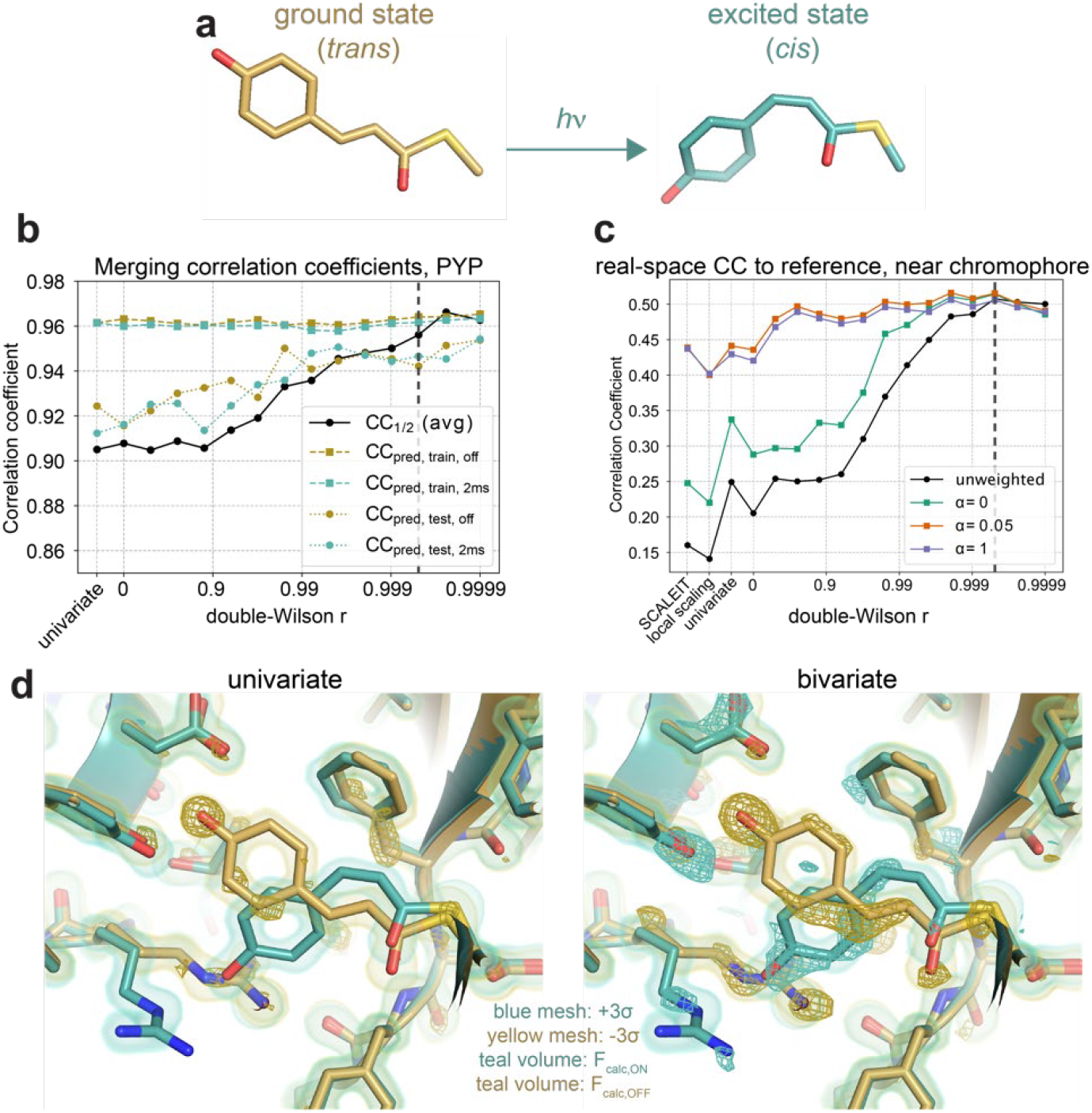
Use of a bivariate prior improves time-resolved difference signal. **a)** The *para*-coumaric acid chromophore of PYP isomerizes when exposed to blue light. **b)** Merging statistics of the 2ms and OFF structure factor amplitudes as a function of *r*. **c)** Correlation coefficient of the weighted (α=0, green; α=0.05, red; α=1, blue) and unweighted (black) observed difference maps with the 2ms−OFF (dark) calculated difference map (*F*_C,2ms_ - *F*_C,OFF_), in a region 10 Å around the chromophore. α weights the contributions of large structure factor differences (see **Methods**). The correlation coefficient is plotted for the univariate prior, with and without applying local scaling or SCALEIT^19^, and as a function of the double-Wilson parameter *r*. Vertical dashed black line in panels **b-d** indicate the *r* value, 0.9995, that parametrizes the bivariate prior for merged data displayed in the following panel. **d) left:** Observed difference map (teal and yellow mesh, contoured to 3σ, unweighted), from merging with a univariate prior. The map is overlaid with models (2ms and dark models colored with teal and yellow sticks, respectively) and the electron density at 2ms after excitation and in the dark state (teal and yellow surfaces, respectively; contoured to 1.5σ), based on calculated structure factors *F*_2ms_ and *F*_off_ from PDB entry 1TSO. **Right:** Observed 2ms−OFF difference map (yellow and teal mesh, contoured to 3σ, unweighted), from merging with a bivariate prior, *r* = 0.9995. F_2ms_ map and F_OFF_ map as in left panel.

It is instructive to compare our scaling approach with commonly used^16,22,42,43^ weights that are heuristic modifications of a weighting scheme proposed in ref^44^, inspired by Bayesian statistics (**Methods**). These weights reduce the contributions of estimated structure factor amplitude differences (Δ*F*) that are large or have large estimated errors (σ(Δ*F*)). A coefficient α, used in calculating the weights, tunes the contribution of large Δ*F* to the difference map (**Methods**). In the case of the PYP data (**Figure 4c**) we find that suppression of large Δ*F*(*α*> 0) is necessary to achieve a similar effect as the bivariate prior. That is, the bivariate prior suppresses spuriously large contributions to the difference map without the need for heuristic weighting schemes.

### Serial Femtosecond Crystallography

Serial femtosecond X-ray crystallography (SFX) of microcrystals probed by intense x-ray free-electron laser (XFEL) pulses has dramatically expanded the range of systems and timescales accessible to time-resolved studies. To determine whether multivariate priors could also improve the accuracy of SFX data processing, we sought to extract anomalous signal from an SFX dataset collected for thermolysin, an enzyme containing a catalytic zinc (Zn) ion in its active site. Specifically, we processed 3,160 images from a much larger dataset (CXIDB 81^45,46^). To account for the serial monochromatic data collection strategy, we included an Ewald offset and per-image layers in our scaling model, as before^22^. We split Bijvoet pairs into two half-datasets before relating these half-datasets by a joint prior during scaling in Careless. The anomalous difference map peak height of the catalytic zinc ion (ZN317) increases from 15σ without a bivariate prior to 28σ for the optimal *r* value (**Figure 5a**). Additionally, as *r* is varied, more associated calcium ions become detectable. Using a univariate prior, only two of four calcium ions appear above 3σ (an approximate noise threshold). After scaling with a multivariate prior, all four previously modeled calcium ions are detectable (**Figure 5a-b**). In prior analyses of a thermolysin dataset with four times as many images, anomalous difference signal has previously been detected at a site adjacent to the bound ZN317^45,47^. When scaling our smaller dataset with a uniform prior, this site was not detectable^22^. Upon scaling with a multivariate prior, however, this site can be clearly identified (**Figure 5b, black arrow**). We note that, again, these anomalous difference maps can also be improved by weighting (**Figure S7**), but that this is substantially less effective than the use of a multivariate prior during scaling. In summary, a bivariate prior strongly improves the detection of anomalous signal from a small SFX dataset.

**Figure 5:**
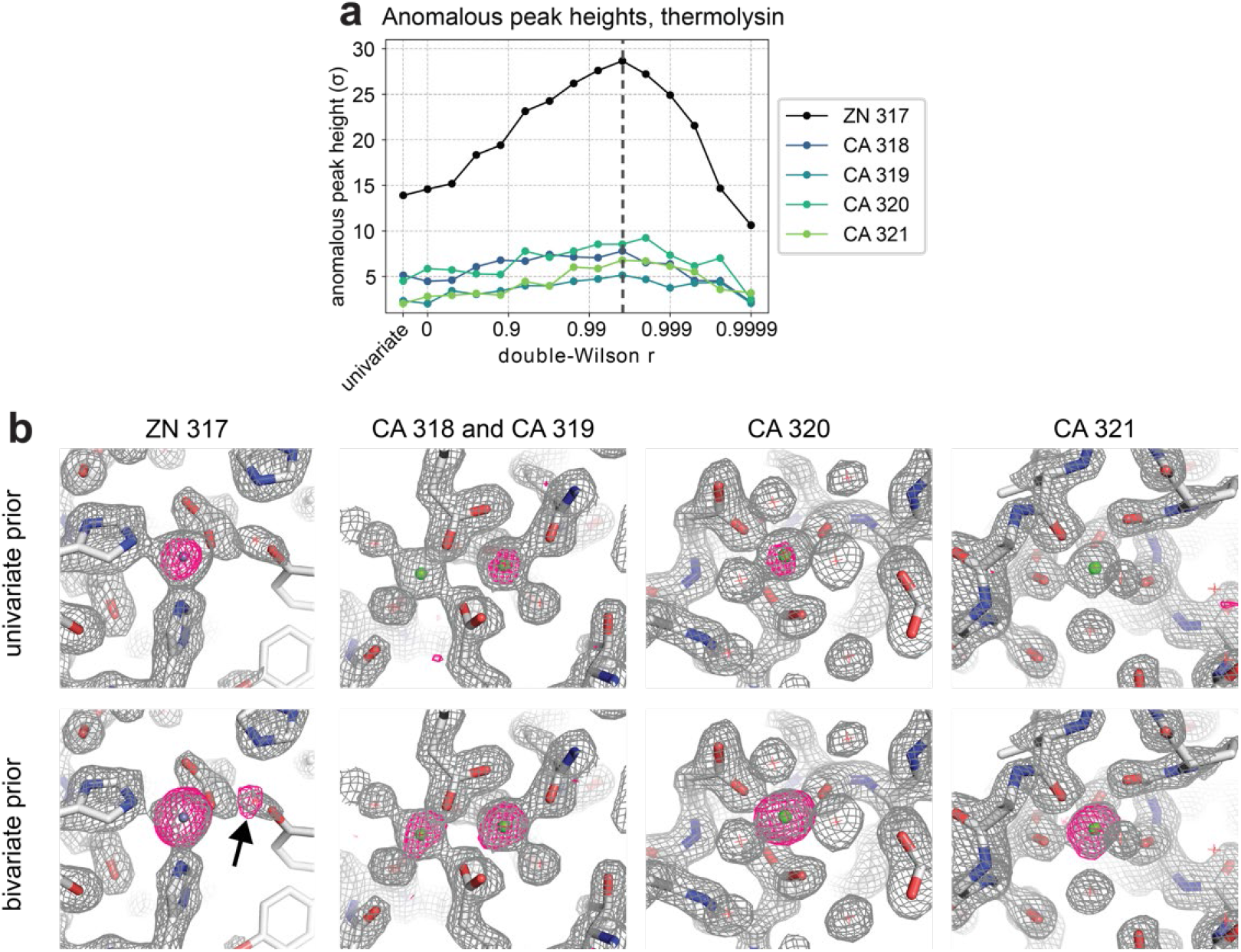
Scaling and merging with a bivariate prior improves the anomalous signal in a serial XFEL experiment. **a)** Anomalous difference map peak heights in thermolysin, shown for zinc and four calcium sites, as a function of the double-Wilson parameter *r*. Vertical dashed black line in panel **a** indicates the *r* value, 0.9961, that parametrizes the bivariate prior for merged data displayed in the following panel. **b) first column:** Region near the thermolysin-bound zinc atom, with observed electron density map (2*mF*_*o*_-D*F*_c_) in gray, contoured at 2σ, and anomalous difference omit map in magenta, contoured at 5σ. Arrow indicates the location of an alternative zinc binding site revealed by merging with the bivariate prior. **Next three columns:** Anomalous difference omit maps of thermolysin calcium sites, carved to 1.5 Å of the model and contoured to 4σ (magenta).

### Fragment screening

Finally, crystallographic drug fragment screens now enable the rapid discovery of pharmacological lead compounds against proteins of interest, e.g., SARS CoV-2 proteins^9,11,13,48^. In these studies, thousands of crystals of a protein of interest are each soaked with a “drug fragment”—a small-molecule building block for larger drug molecules. To determine if joint scaling can improve the detection of bound fragments in difference maps, we analyzed data from a fragment screening experiment totaling one *apo* dataset and sixteen *holo* datasets identified by PanDDA^49^. This dataset contains structures of SARS-CoV2 nonstructural protein 3 macromolecular domain 1 (Nsp3 Mac1) with ligands from a fragment screening library bound to the Mac1 adenosine-binding site. We ran *Careless* on unmerged intensities, adding several metadata keys including one-hot encodings of each dataset (**Methods**). By one-hot encoding, we allow the model to express differences in scale between fragment-bound datasets.

Additionally, we set each *holo* dataset to depend on the *apo* dataset through a joint prior, while assuming conditional independence between *holo* datasets (**Figure 2b**). We find that difference density maps visually improve upon scaling with a multivariate prior (**Figure 6a, Figure S8**). We tracked the largest ligand peaks in the difference map for a given dataset while we varied *r*. We find that in all datasets, this signal is strongest when scaling with a multivariate prior (**Figure 6b**). The *r* value for the tallest peak differs for each *holo* dataset. Without special modifications to scaling and merging, we obtain peak heights on par with those produced by PanDDA, which relies on real-space post-processing (**Table S2**). To determine if the *apo*-*holo* dataset structure factor correlation, measured by *r*, relates to ligand occupancy, we examined the relationship between optimal *r* and the PanDDA background density correction factor (BDC, a parameter which scales inversely with ligand occupancy and accounts for crystal-to-crystal variation in soaking and data quality)^12^. We find a moderate correlation with *r* (Pearson *r* = 0.505, *p* < 0.05) (**Figure S9a**). Merging statistics also improve when scaling with a bivariate prior (**Figure S9b**): in particular, the gap between the training and test *CC*_pred_ is reduced for higher *r*, indicating that the bivariate prior reduces overfitting of the scales.

**Figure 6.**
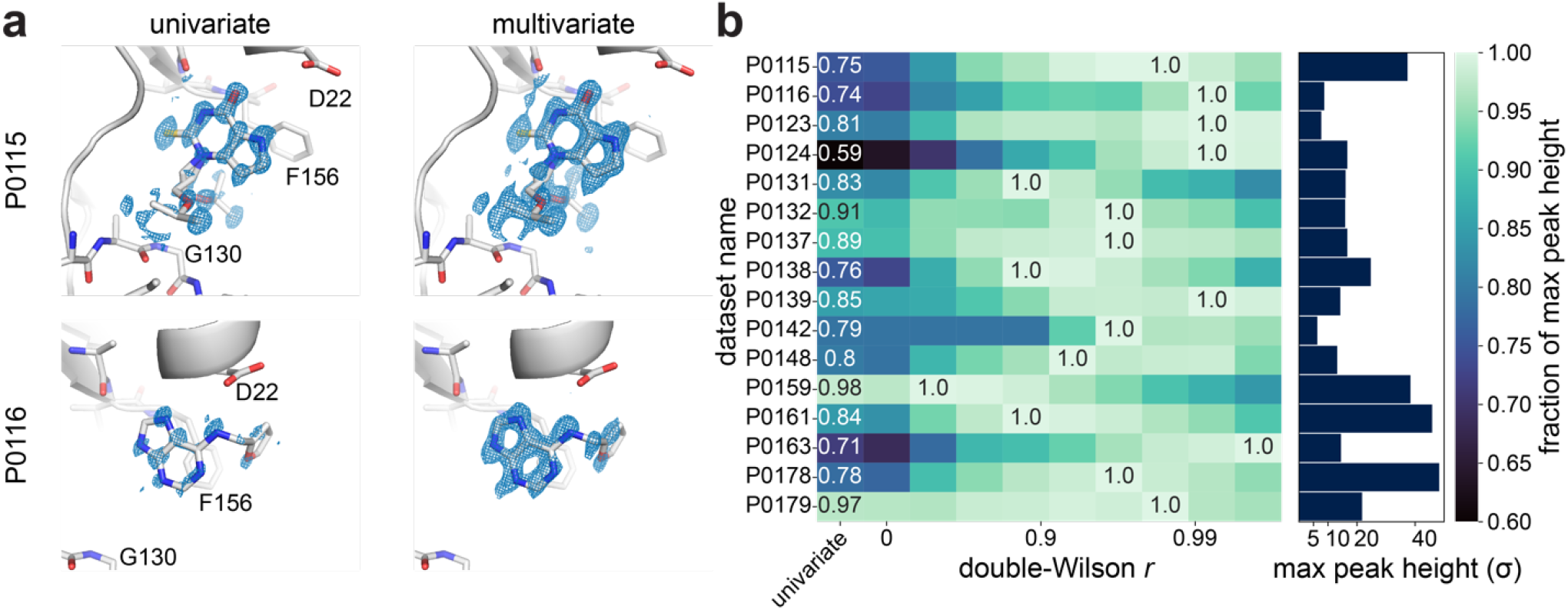
Use of a multivariate prior improves bound fragment signal in a drug fragment screen. **a)** *holo* − *apo* isomorphous difference maps from fragment screening of Mac1, scaled with a univariate or multivariate prior. All maps carved to 1.5 Å of the ligand and contoured to +3σ. **b)** Tallest positive peak in each *holo* − *apo* difference map as a function of the double-Wilson *r* parameter. For a given *holo* dataset, the tallest positive peak is measured. The maximum peak height of that peak, across all double-Wilson *r* values, is normalized to 1.0. The maximum peak height is labeled and reported in a bar chart (**right**). Also labeled is the fraction of the maximum peak height for the tallest peak in the univariate datasets (**left**). The r value, for each dataset’s maximum peak height, parametrizes the bivariate prior for merging the electron density displayed in panel **a**.

This example shows that the difference map quality improves when scaling high-resolution monochromatic data with a bivariate prior. It would be natural to expect that the benefits of a multivariate prior during scaling would keep increasing with the number of datasets included. At present, this is not the case, likely because we do not account for crystal-to-crystal variation in the average unit cell.

## Discussion

Bayesian statistics have been applied extensively in macromolecular crystallography, playing an important role in solutions to the phase problem^28^ and maximum-likelihood-based refinement^50^. Here, we establish that a Bayesian formalism enables the sensitive detection of subtle chemical and time-dependent signals and ligand binding from comparative crystallographic measurements by introducing a multivariate prior describing the anticipated correlations among related datasets. Although this formalism proves effective even in the simple form presented here, there are several promising extensions to further improve the detection of structural change. For example, high-quality reference datasets are commonly measured for serial time-resolved crystallography experiments, but these data are not commonly used to improve scaling. Such datasets could be readily added to the Bayesian networks shown in **Figure 2** to better condition scaling. Second, we currently treat the correlation parameter *r* as a fixed, global hyper-parameter, yet correlations may vary between pairs of datasets (e.g., as a function of ligand occupancy) and vary with resolution. These correlations could be estimated from the data. Third, as not every node in **Figure 2** needs to be observed, our approach allows for interpolation of missing observations.

The effects of perturbations in comparative crystallography, whether due to laser excitation^51^, temperature^15,52^, electric field^16^, or small-molecule binding^11^, often only affect a fraction of the molecules in the crystal. As an important consequence, the resulting mixed structure factor amplitudes tend to be slightly smaller than the ground-state (OFF) amplitudes^53^, and the resulting ON-OFF difference maps tend to resemble a negative image of the OFF electron density. It is common practice to rescale scaled data to place OFF and ON structure factor amplitudes on a common scale (as we do, implicitly, in the present work). In the developed statistical framework, correlation of an isomorphous difference map with the OFF electron density, the true ON-OFF difference map, and the map error can all be calculated exactly (**Supplementary Notebook 6**), supporting the common observation that correlation with the OFF map can be removed with minimal effect on difference map quality (**Figure S12**).

Finally, we observe that the developed framework also implies a joint distribution of the unobserved crystallographic phases across related datasets (**SI** and **Figure S13**). As the traditional crystallographic data processing paradigm begins to give way to machine learning methods, we anticipate that these implied correlations can play a key role in learning crystallographic movies directly and efficiently from time-resolved diffraction data.

## Methods

### Implementation details

The multivariate Wilson distribution we present here is implemented as a prior distribution in the Careless software package. Careless uses gradient-based optimization to maximize a Bayesian objective function^54,55^ which is the sum of the expected log likelihood of the observed data plus the Kullback-Leibler divergence between the structure factors and the prior distribution. The likelihood of the data describes the probability of the observed diffraction intensities, *I*_*h,i*_, conditional on scale Σ_*h,i*_, and structure factor amplitudes *F*_*h*_. The likelihood also takes into account the estimated errors 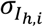 in the observed intensities. The objective function is computed as described in equation 21 and 22 of Dalton et al.^22^ using reparameterized samples^56^ from a truncated normal surrogate posterior (variational distribution). For clarity, we reproduce this objective function,

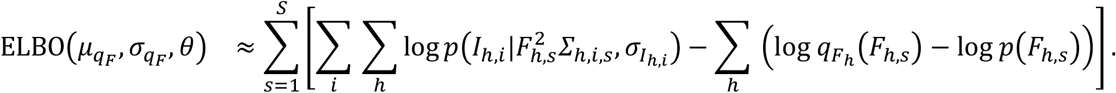

The Evidence Lower BOund (ELBO) depends on the parameters of the surrogate posterior distributions, *q*_*F*_ and *q*_*∑*_. *q*_*∑*_ is parameterized by a neural network with parameters *θ. q*_*F*_ is parameterized by independent truncated normal distributions for each reflection with location and scale parameters {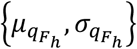}supported on [0, ∞) for centric reflections and (0, ∞) for acentrics. *I*_*h,i*_ refers to the intensity of a particular observation *i* for reflection *h. s* indexes reparameterized samples from the surrogate posteriors,

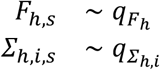

Reparameterization is implemented in the TensorFlow Probability library^57^. The prior distribution enters the objective function through the ultimate term, log *p*(*F*_*h,s*_).Previously,^22^ we exclusively used a univariate Wilson distribution as a prior for this term,

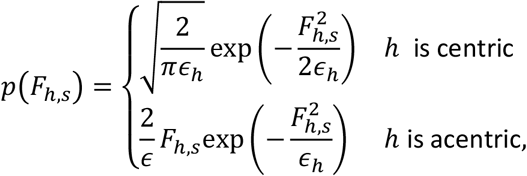

where *ϵ*_*h*_ is the multiplicity of the reflection *h*, an integer value determined by the crystal’s symmetry. Note that because the scale function, Σ_*h*_, is parametrized separately within Careless, it does not appear in expressions for *p*(*F*_*h,s*_).

One of the features of variational inference is its flexibility. In this work we leverage the inherent flexibility of reparameterization-based variational inference to extend the Careless package with a multivariate prior distribution. To do so, the Careless command line interface requires users to supply separate unmerged data sets for each node they wish to model (as in **Figure 2**). Careless asks the user to specify the parent of each node and the expected double-Wilson parameter *r*. Internally, this changes how the prior is calculated as follows,

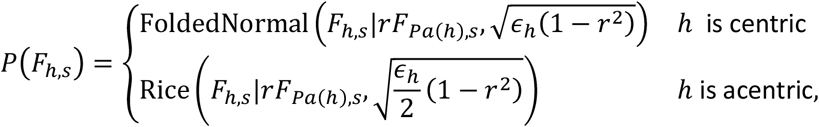

where *Pa* (*h*) denotes the parent reflection of *h*. Where a particular reflection has no parent because it is the root node, the resolution range of parent and child differ, or the parent is systematically absent, the prior defaults to the univariate Wilson model. The folded normal distribution’s probability density function is

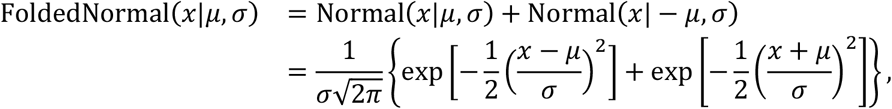

while that of the Rice distribution is

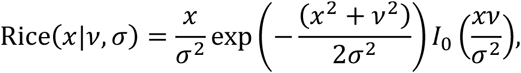

where *I*_0_ denotes the modified Bessel function of the first kind with order zero. For numerical stability, both probability densities are computed directly in log space.

### PYP

Collection of the Laue diffraction data for photoactive yellow protein was described previously^58^. Indexing, geometry refinement, wavelength assignment, and integration were performed in Precognition v5.2.2. Precognition integrated intensities were written to mtz files and then scaled and merged with Careless v0.4.1, with use of a bivariate prior on the OFF and 2ms timepoint mtz files. Fifteen different double-Wilson *r* values were used, with *r* varied for each Careless run as 1 − 0.5^run number^. Correlation coefficients were computed using *careless*.*cchalf, careless*.*ccpred*, and custom scripts based on *gemmi*^59^ and *reciprocalspaceship*^60^. Isomorphous difference maps were first phased with an OFF model refined in Phenix 1.20.1^61^ from PDB ID 2PHY^62^, then weighted using *reciprocalspaceship* according to

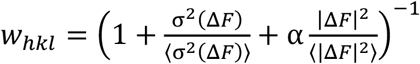

for structure factor amplitude differences Δ*F* (short for Δ*F*_*hkl*_) and their errors σ (Δ*F*), with angled brackets indicating averages over Miller indices. Local scaling was performed using a custom script available at (https://github.com/Hekstra-Lab/dw) and SCALEIT (CCP4 software suite version 7.1) was run from *rs-booster* version 0.1.2 (https://github.com/rs-station/rs-booster) as *rs*.*scaleit* (**Supplementary Information, “Partial Excitation”**).

Difference maps were visualized in *PyMOL*^63^. Data processing scripts can be found in the Zenodo deposition.

### Lysozyme data collection

NaI-soaked lysozyme crystals were prepared as described previously^64^. Monochromatic data were collected on the Northeastern Collaborative Access Team (NE-CAT) beamline 24-ID-C (Advanced Photon Source, Argonne National Laboratory) on 12 August 2020. Diffraction data were collected at ambient temperature (about 295 K). Three 1440-image passes (720°) were collected from a single crystal of lysozyme with an exposure time of 0.1 s and an oscillation angle of 0.5°. The incident X-ray intensity, at an energy of 11.95 keV, was attenuated to 0.5% transmission, which corresponds to an estimated flux of 2.4 × 10^10^ photons s^−1^. Data were collected using helical acquisition, so that dose was evenly distributed along the crystal. The PILATUS 6M-F detector (Dectris) was positioned at the minimal distance of 150 mm.

Laue data were collected on the same day at BioCARS (Advanced Photon Source, Argonne National Laboratory), similar to data collection described in previous work^16^, except that Laue stills were collected in 1° steps, for a total of 3,000 images. Data were collected at ambient temperature (about 295 K).

### Thermolysin and lysozyme

Careless 0.4.1 was run with a bivariate prior, sweeping over fifteen double-Wilson *r* values on two datasets, one of thermolysin containing 3160 images collected from a serial XFEL experiment, CXIDB81^45^, and one of lysozyme containing the first 999 images from the above data collection. Lysozyme data were indexed and integrated using *laue-dials* (https://github.com/rs-station/laue-dials; manuscript in preparation). Thermolysin unmerged intensities (in DIALS .*pickle* format) were obtained from CXIDB 81. Both datasets were split according to Friedel symmetry and scaled and merged in Careless, applying the bivariate prior to relate anomalous half-datasets. Next, the anomalous half-datasets were combined into single datasets of merged structure factor amplitudes.

Correlation coefficients on these datasets were computed using *careless*.*cchalf, careless*.*ccpred*, and *careless*.*ccanom*^36^. Phenix version 1.20.1^61^ was used to phase the resultant structure factor amplitudes by isomorphous replacement with PDBID 2TLI in the case of thermolysin and with a high-resolution monochromatic model in the case of lysozyme (**Table S1**). Additionally, an anomalous omit map was phased. The phases and omit map were then used for visualization of anomalous omit peaks with *PyMOL* and measurement of peak heights with *reciprocalspaceship*. Data processing scripts can be found in the Zenodo deposition.

### Fragment screening of Mac1

Data collection and analysis of a crystallographic drug fragment screen of SARS-CoV-2 non-structural protein 3 (Nsp3) macrodomain 1 (Mac1) have been previously described^11^. For the work described here, one unmerged *apo* dataset and sixteen unmerged *holo* datasets HKL files were converted to MTZ files and then processed with Careless 0.4.1, with use of a univariate or multivariate prior to relate the OFF and sixteen holo files. The double-Wilson *r* was set to a uniform value for all sixteen *apo*-*holo* dependencies, and was varied between Careless runs as 1 − 0.5^run number^. Additionally, we introduced a control *apo* dataset, a duplicate of the reference that also depended on the reference *apo* dataset, and difference maps were computed by subtracting each *holo* dataset from the control *apo* dataset. To compensate for the duplication of the reference *apo* dataset, we multiply the errors of the intensities by 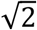. Metadata keys included observed Miller indices, detector position and positional encoding, image number, the azimuthal angle PSI^65^, diffracted beam direction parameters ALF1 and BET1^66^, and dataset one-hot encoding. Correlation coefficients were computed using *careless*.*ccpred. Apo* phases were obtained by refinement of a model against the *apo* mtz scaled using a univariate prior. Refinement was performed using PHENIX v1.20.1. The *apo* phases were then used for creating isomorphous difference maps of each *holo* minus *apo* dataset. We created difference maps and calculated peak heights using *rs-booster* (https://github.com/rs-station/rs-booster). PanDDA 1-background density correction (1-BDC) values were taken from an analysis that included all previously-reported datasets^11,49^. Data processing scripts can be found in the Zenodo deposition.

## Data Availability

Supplementary notebooks, including analyses used for generating **Figures 1-2** and **S1-4,10-13** are available in a GitHub repository at https://github.com/Hekstra-Lab/dw. The monochromatic sodium-iodide-soaked lysozyme structure presented here has been deposited in the Protein Data Bank^67^ with PDB ID 9B7C. All merged structure factors, crossvalidation data, and analysis used to generate **Figures 3-6** and **S5-9** have been deposited in Zenodo under accession code 10.5281/zenodo.11099739 at https://doi.org/10.5281/zenodo.11099739. The analyses, without merged structure factors, are included in a GitHub repository at https://github.com/Hekstra-Lab/dw-examples.

## Code Availability

The source code used for generating all figures and tables are available in the GitHub repositories https://github.com/Hekstra-Lab/dw and https://github.com/Hekstra-Lab/dw-examples as well as deposited in Zenodo at https://doi.org/10.5281/zenodo.11099739. The multivariate prior scaling model is implemented in the Careless python package, available in the GitHub repository https://github.com/rs-station/careless.

## Acknowledgements

We thank Dr. Michael Socolich (University of Chicago) for sodium iodide-soaked lysozyme crystals; Drs. Vukica Šrajer (University of Chicago) and Marius Schmidt (University of Winconsin) for providing Laue time-resolved data, Drs. Galen Correy and James S. Fraser (UCSF) for unmerged Mac1 drug fragment screening data, and Drs. TJ Lane, Vukica Šrajer, and Randy Read for comments on the manuscript. We thank the staff at the Northeastern Collaborative Access Team (NE-CAT), beamline 24-ID-C of the Advanced Photon Source (APS) for assistance with data collection, in particular Dr. Igor Kourinov. We thank the staff at the BioCARS facility at beamline 14-ID-B of APS, in particular Drs. Robert Henning and In-Sik Kim, for assistance with polychromatic data collection. We acknowledge the open-source software projects used in this work including DIALS^68^ GEMMI^59^, Matplotlib^69^, Numpy^70^, Pandas^71^, reciprocalspaceship^60^, SciPy^72^, Seaborn^73^, TensorFlow^74^, TensorFlow Probability^57^, and Careless. This work was supported by the National Institutes of Health (NIH) Director’s New Innovator Award (DP2-GM141000) to D.R.H. H.K.W. was supported by the National Science Foundation Graduate Research Fellowship under Grant No. DGE2140743. K.M.D. holds a Career Award at the Scientific Interface from the Burroughs Wellcome Fund. NE-CAT beamlines are supported by the National Institute of General Medical Sciences (NIGMS), NIH (P30 GM124165). BioCARS is supported by NIGMS under grant number P41 GM118217. The time-resolved set-up at Sector 14 was funded in part through a collaboration with Philip Anfinrud (NIH/National Institute of Diabetes and Digestive and Kidney Diseases). APS, a U.S. Department of Energy (DOE) Office of Science User Facility is operated for the DOE Office of Science by Argonne National Laboratory under Contract No. DE-AC02-06CH11357. This publication’s contents are solely the responsibility of the authors and do not necessarily represent the official views of NIGMS, the NIH, or the DOE.

## Author contributions

D.R.H. and K.M.D. conceived the project. D.R.H. developed the theory and K.M.D. implemented the algorithm. J.B.G. and K.M.D. collected lysozyme anomalous diffraction data. D.R.H., H.K.W., and M.A.K. analyzed the data. D.R.H., H.K.W., and K.M.D. drafted the manuscript with input and edits from all authors.

