## Supplementary Information for "Sensitive Detection of Structural Differences using a Statistical Framework for Comparative Crystallography"

#### S1. A statistical model for correlations of related structure factors

##### The Wilson distribution

The Wilson distribution provides a simple, long-standing model for the statistical distribution of the structure factors of macromolecular crystals[1,2]. Briefly, structure factors can be calculated as a sum over contributing atoms, $j$, in the complex plane, that is,

$$\boldsymbol{F}_h = \sum_j f_{\boldsymbol{h}j}^B e^{2\pi i(\boldsymbol{h}^T \boldsymbol{x}_j)} \qquad [1]$$

where $f_{\boldsymbol{h}j}^B$ captures the finite size of atoms and their thermal disorder, and $\boldsymbol{h}^T\boldsymbol{x}_{\boldsymbol{j}}$ is the dot product of the atomic fractional coordinate vector $\boldsymbol{x}_{\boldsymbol{j}}$ and Miller indices $\boldsymbol{h}$. (We will use boldface notation for vectors and the complex structure factors $\boldsymbol{F}_h$, while reserving italics for structure factor amplitudes $F_h \equiv |\boldsymbol{F}_h|$ and other scalar quantities.) For most reflections, as each subsequent atom is added to the calculation, the growing structure factor performs a nearly random walk in the 2D complex plane, and the central limit theorem can be applied[2]. The rest of the reflections are centric and thus constrained to lie on a line through the origin due to crystallographic symmetry restrictions[2]. Thus, the acentric structure factors approximately follow a bivariate normal, while centrics follow a univariate normal distribution, both with zero mean. To illustrate, for the real and imaginary components of an acentric structure factor $Re(\boldsymbol{F}_{\boldsymbol{h}})$ and $Im(\boldsymbol{F}_{\boldsymbol{h}})$ respectively,

$$P(Re(\boldsymbol{F}_{\boldsymbol{h}}), Im(\boldsymbol{F}_{\boldsymbol{h}})) = \mathcal{N}\left(0, \frac{1}{2}\epsilon_{\boldsymbol{h}}\Sigma_{\boldsymbol{h}} I\right) \qquad [2]$$

where the multiplicity $\epsilon_h$ accounts for crystallographic symmetry[2], $I$ is a 2x2 identity matrix, and $\Sigma_{\boldsymbol{h}}$ is an overall scale. In the Wilson model, $\Sigma_{\boldsymbol{h}}$ represents the total scattering power of atoms in the unit cell as a function of resolution (that is: $\mathbb{E}(|\boldsymbol{F}_{\boldsymbol{h}}|^2) = \epsilon_{\boldsymbol{h}}\Sigma_{\boldsymbol{h}} = Re(\boldsymbol{F}_{\boldsymbol{h}})^2 + Im(\boldsymbol{F}_h)^2$. It will be convenient to normalize structure factors $\boldsymbol{E}_{\boldsymbol{h}} = \frac{\boldsymbol{F}_{\boldsymbol{h}}}{\sqrt{\epsilon_{\boldsymbol{h}}\Sigma_{\boldsymbol{h}}}}$, such that

$$P(Re(\boldsymbol{E}_{\boldsymbol{h}}), Im(\boldsymbol{E}_{\boldsymbol{h}})) \sim \mathcal{N}\left(0, \frac{1}{2} I\right) \quad \text{for acentric structure factors and} \qquad [3a]$$

$$\boldsymbol{E_h} \sim \mathcal{N}(0,1) \qquad \text{for centric structure factors.} \qquad [3b]$$

For initial inspection of already-merged data, we will treat $\Sigma$ as an empirical normalization constant (**Supplementary Notebooks 1-3,6**). When scaling and merging diffraction data, we note that *Careless* treats $\Sigma$ as the scale function that needs to be inferred.

In X-ray crystallography the phase of the structure factor is not observed[3], so the distribution of the amplitudes of structure factors (here denoted $E_{\boldsymbol{h}}$) is of great interest. Following Wilson[2], this distribution can be obtained by first converting from Cartesian to polar coordinates and then integrating over the unknown phases to yield the standard Wilson distribution,

$$P(E_{\boldsymbol{h}}) = \begin{cases} \sqrt{\frac{2}{\pi}} \exp\left(-\frac{1}{2} E_{\boldsymbol{h}}^2\right) \text{ for } \boldsymbol{h} \text{ centric} \\ 2E_{\boldsymbol{h}} \exp(-E_{\boldsymbol{h}}^2) \quad \text{ for } \boldsymbol{h} \text{ acentric} \end{cases} \qquad [4]$$

This distribution is the basis for many models of structure factor amplitudes[1].

#### The bivariate Wilson distribution

When considering a pair of data sets collected under similar conditions, we treat the sum in eq. [1] for two data sets as correlated random walks[4] with correlation coefficients, $r$, between the corresponding real components, and likewise between imaginary components (**Figure 1a**). In this approach, the joint probability distribution of two acentric structure factors $\boldsymbol{E}_{\boldsymbol{h}}^A$ and $\boldsymbol{E}_{\boldsymbol{h}}^B$ is[5]:

$$P\left(Re(\boldsymbol{E}_{\boldsymbol{h}}^A), Im(\boldsymbol{E}_{\boldsymbol{h}}^A), Re(\boldsymbol{E}_{\boldsymbol{h}}^B), Im(\boldsymbol{E}_{\boldsymbol{h}}^B)\right) = \mathcal{N}\left(0, \frac{1}{2}\begin{bmatrix} 1 & 0 & r & 0 \\ 0 & 1 & 0 & r \\ r & 0 & 1 & 0 \\ 0 & r & 0 & 1 \end{bmatrix}\right) \qquad [5]$$

with the parameter $r$ henceforth called the "double-Wilson $r$". The conditional probability distribution of $\boldsymbol{E}_{\boldsymbol{h}}^B$ given $\boldsymbol{E}_{\boldsymbol{h}}^A$ can now be calculated as $P\left(Re(\boldsymbol{E}_{\boldsymbol{h}}^B), Im(\boldsymbol{E}_{\boldsymbol{h}}^B) | \boldsymbol{E}_{\boldsymbol{h}}^A\right) = \mathcal{N}\left(r\boldsymbol{E}_{\boldsymbol{h}}^A, \frac{1}{2}(1 - r^2)I\right)$, and the conditional probability distribution of their amplitudes (i.e. $E_{\boldsymbol{h}}^B$ conditional on $E_{\boldsymbol{h}}^A$) can be calculated by marginalizing over both the unknown phase of $\boldsymbol{E}_{\boldsymbol{h}}^A$ and the unknown phase difference between $\boldsymbol{E}_{\boldsymbol{h}}^A$ and $\boldsymbol{E}_{\boldsymbol{h}}^B$[6], yielding the Rice distribution,

$$P(E_h^B) \sim Rice(\nu, \sigma^2) = \frac{x}{\sigma^2} \exp\left(\frac{-(x^2+\nu^2)}{2\sigma^2}\right) I_0\left(\frac{x\nu}{\sigma^2}\right)$$

With $x = E_h^B$ and parameters $\nu = rE_{\boldsymbol{h}}^A$ and $\sigma^2 = \frac{1}{2}(1 - r^2)$ and $I_0$ the modified Bessel function of the first kind with order zero. This derivation is elaborated upon in **Supplementary Notebook 5**.

Now for centric structure factors $\boldsymbol{E}_{\boldsymbol{h}}^A$ and $\boldsymbol{E}_{\boldsymbol{h}}^B$ we have:

$$P(\boldsymbol{E}_h^A, \boldsymbol{E}_h^B) = \mathcal{N}\left(0, \frac{1}{2}\begin{bmatrix} 1 & r \\ r & 1 \end{bmatrix}\right)$$

so that the probability distribution of their amplitudes $E_{\boldsymbol{h}}^B$ conditional on $E_{\boldsymbol{h}}^A$ is

$$P(E_{\boldsymbol{h}}^B | E_{\boldsymbol{h}}^A) = \mathrm{FoldedNormal}(\mu, \sigma^2) = \frac{1}{\sqrt{2\pi\sigma^2}}\left(\exp\left(-\frac{(x-\mu)^2}{2\sigma^2}\right) + \exp\left(-\frac{(x+\mu)^2}{2\sigma^2}\right)\right)$$

for $x = E_h^B$, the magnitude of $\boldsymbol{E}_{\boldsymbol{h}}^{\boldsymbol{B}}$, and parameters $\mu = rE_{\boldsymbol{h}}^A$ and $\sigma^2 = (1 - r^2)$.

Following ref. [5], we note that $r$ often depends on resolution, consistent with a model proposed by Luzzati, such that $r \approx ae^{-bs^2}$, where s is 1/resolution. We also note that $r$ plays a role analogous to $\sigma_A$ in crystallographic refinement[4,7] and can be estimated similarly. To illustrate this, we fit related pairs of structure factor amplitudes to this model, obtaining estimates of $a$ and $b$, in "**Estimation of r from merged structure factor amplitudes"** and in **Supplementary Notebook 3**. We observe that a pair of synthetic datasets obtained by resampling of example observed data (while imposing $r = ae^{-bs^2}$) yields a resolution-dependent correlation (blue line in **Figure S10**) similar to the observed correlation (green line in **Figure S10**). The inferred true correlations between structure factor amplitudes (dashed magenta line in **Figure S10**) is of similar magnitude, indicating most of the decay in correlation with resolution is real rather than due to measurement errors (see also **Supplementary Notebook 2b**).

#### The multivariate Wilson distribution

The conditional independence graphs in Figure 2 all take the form of a tree (a connected, acyclic graph). Given this, it is straightforward to calculate the full covariance matrix for the joint distribution of complex structure factors from the diagram[8] (**Supplementary Notebook 7**). More importantly, we can factorize the joint probability of structure factors as

$$P\left(\boldsymbol{E}_{\boldsymbol{h}}^{(1)}, \boldsymbol{E}_{\boldsymbol{h}}^{(2)}, \boldsymbol{E}_{\boldsymbol{h}}^{(3)}, \ldots\right) = P\left(\boldsymbol{E}_{\boldsymbol{h}}^{(0)}\right) \cdot \prod_{j>0} P\left(\boldsymbol{E}_{\boldsymbol{h}}^{(j)} \middle| \boldsymbol{E}_{\boldsymbol{h}}^{\mathrm{Pa}(j)}\right) \quad [6a]$$

where we set the index of the root of the tree to 0, and $\mathrm{Pa}(j)$ is the "parent node" of node $j$. Whenever this is possible, we can integrate over phase differences factor by factor, and obtain an analogous expression for structure factor amplitudes,

$$P\left(E_{\boldsymbol{h}}^{(1)}, E_{\boldsymbol{h}}^{(2)}, E_{\boldsymbol{h}}^{(3)}, \ldots\right) = P\left(E_{\boldsymbol{h}}^{(0)}\right) \cdot \prod_{j>0} P\left(E_{\boldsymbol{h}}^{(j)} \middle| E_{\boldsymbol{h}}^{Pa(j)}\right). \quad [6b]$$

For acentric reflections, $P\left(E_{\boldsymbol{h}}^{(j)} \middle| E_{\boldsymbol{h}}^{Pa(j)}\right)$ again follows a Rice distribution, now with $\nu = r_j E_{\boldsymbol{h}}^{Pa(j)}$ and $\sigma^2 = \frac{1}{2}\left(1 - r_j^2\right)$, with $r_j$ as the double-Wilson $r$ for the correlation between node $j$ and its parent. For centric reflections, $P\left(E_{\boldsymbol{h}}^{(j)} \middle| E_{\boldsymbol{h}}^{Pa(j)}\right)$ again follows a folded normal distribution with $\mu = r_j E_{\boldsymbol{h}}^{Pa(j)}$ and $\sigma^2 = \left(1 - r_j^2\right)$. We provide a numerical demonstration of equation [6b] in **Supplementary Notebook 7**. We will refer to the distribution in eq. [6b] as the *multivariate Wilson distribution*.

#### Estimation of *r* from merged structure factor amplitudes

The double-Wilson parameter *r* is closely related to the correlation between two datasets. In **Supplementary Notebook 2a**, we find numerically that the Pearson correlation between two synthetic datasets of normalized structure factor amplitudes, $E_1$ and $E_2$, is approximately $r^2$. Indeed, within the multivariate Wilson model, the *squares* of the structure factor amplitudes are correlated exactly as $r^2$ (see ref. [9]). Using the former relation, we may obtain a resolution-independent value for the double-Wilson r, but we may also fit the double-Wilson *r* as a function of resolution. To do so, we model variation in the amplitudes as follows. We model $E_{1,obs} = x + \eta_1$, where $x$ is the true value of $E_1$ and $\eta_1$ is measurement error. We model the second dataset as $E_{2,obs} = x + \epsilon + \eta_2$, where $\epsilon$ is the true difference between $E_2$ and $E_1$. We assume that $x, \epsilon, \eta_1$ and $\eta_2$ are uncorrelated with variances $\sigma_x^2$, $\sigma_\epsilon^2$, $\sigma_1^2$ and $\sigma_2^2$. Then, we have

$$\rho_{obs} = \rho(x + \eta_1, x + \epsilon + \eta_2) = \frac{\sigma_x^2}{\sqrt{(\sigma_x^2 + \sigma_1^2)(\sigma_x^2 + \sigma_\epsilon^2 + \sigma_2^2)}}$$

and

$$\rho_{true} = \rho(x, x + \epsilon) = \frac{\sigma_x^2}{\sqrt{(\sigma_x^2)(\sigma_x^2 + \sigma_\epsilon^2)}}$$

so that

$$\rho_{obs}^{-2} = \rho_{true}^{-2} + \frac{\sigma_1^2 + \sigma_2^2}{\sigma_x^2} + \frac{\sigma_1^2 \sigma_2^2}{\sigma_x^4} + \frac{\sigma_1 \sigma_2}{\sigma_x^2}(\rho_{true}^{-2} - 1)$$

$\sigma_x^2$ is a known property of the Wilson distribution, allowing us to relate the true correlation between $E_1$ and $E_2$ to the observed correlation in the presence of measurement errors. With this relationship in hand, we can then find the value of $r$ by approximating $\rho_{true} \approx r^2$ and, if modeling resolution-dependence, of $a, b$ such that $r = a \cdot e^{-bs^2}$.

#### Normalizing structure factors

The normalized structure factor amplitudes used for the examples in **Figures 1 and S1-4** were obtained from datasets that were processed using DENZO/SCALEPACK[10], XSCALE[11], or SCALA[12]. The structure factor amplitudes $F_h$ produced by these software are not normalized. Additionally, these structure factors may contain residual systematic errors in scaling. To assess the correspondence of these data to our statistical model, we first needed to normalize the $F_{\boldsymbol{h}}$. To do so, we introduced an ad hoc normalization procedure (which is not relevant to the results in **Figures 3-6**, which were obtained by scaling and merging using Careless). To this end, we first observe that the Wilson model can be formulated more generally as

$$P(F_{\boldsymbol{h}}) = \begin{cases} \sqrt{\frac{2}{\pi\epsilon_{\boldsymbol{h}}\Sigma_{\boldsymbol{h}}}} \exp\left(-\frac{F_{\boldsymbol{h}}^2}{2\epsilon_{\boldsymbol{h}}\Sigma_{\boldsymbol{h}}}\right) \text{for } \boldsymbol{h} \text{ centric} \\ 2\frac{F_{\boldsymbol{h}}}{\epsilon_{\boldsymbol{h}}\Sigma_{\boldsymbol{h}}} \exp\left(-\frac{F_{\boldsymbol{h}}^2}{\epsilon_{\boldsymbol{h}}\Sigma_{\boldsymbol{h}}}\right) \quad \text{for } \boldsymbol{h} \text{ acentric} \end{cases}$$

where $\epsilon_{\boldsymbol{h}}$ is the multiplicity of reflections, and $\Sigma_{\boldsymbol{h}}$ dictates the mean square intensity of reflections at (or nearby) Miller index $\boldsymbol{h}$ (short for $(h, k, l)$). To use Wilson statistics, we must learn how $\Sigma$ varies across reciprocal space—or, equivalently, infer normalized structure factor amplitudes $E_{\boldsymbol{h}} = \frac{F_{\boldsymbol{h}}}{\sqrt{\epsilon_{\boldsymbol{h}}\Sigma_{\boldsymbol{h}}}}$, which obey $\langle E^2 \rangle = 1$ and follow the Wilson distribution for normalized structure factors.

In **Supplementary Notebook 1**, we normalize structure factors in three steps. First, we find an optimal anisotropic *B* matrix that best approximates a standard Wilson distribution. That is, we model $\Sigma_{\boldsymbol{h}}$ as $\Sigma'_{\boldsymbol{h}} = a \cdot \exp\left(-\frac{1}{2} B s_{\boldsymbol{h}}^2\right)$, with $s_{\boldsymbol{h}} = \frac{1}{d_{hkl}}$, and find the maximum of the likelihood function $l(a, B) = P(\{F_{\boldsymbol{h}}\} \mid a, B) = \prod_{\boldsymbol{h}} P(F_{\boldsymbol{h}} \mid a, B)$. Such likelihood maximization under the Wilson distribution has been previously described[13,14].

At this point, the mean squared intensity is often still not uniform across reciprocal space. This is for several reasons, such as incomplete correction for absorption artifacts during scaling, as well as the occurrence of regularities in protein structure (e.g. secondary structure). A heuristic to correct for this is to propose that $\Sigma_{\boldsymbol{h}}$ be modulated by position in reciprocal space as:

$$\Sigma''_{\boldsymbol{h}} = \left( \sum_{\boldsymbol{n}} A_{\boldsymbol{n}} \cos\left(\frac{2\pi \boldsymbol{h}^T \boldsymbol{n}}{L}\right) + B_{\boldsymbol{n}} \sin\left(\frac{2\pi \boldsymbol{h}^T \boldsymbol{n}}{L}\right) \right) \Sigma'_{\boldsymbol{h}}$$

with the anisotropically determined value $\Sigma'_{\boldsymbol{h}}$, the Fourier component $\boldsymbol{n} \in [0, 1, 2, 3, 4]^{\boldsymbol{3}}$, and the entire sum computed in a box with side lengths $L$ that bounds the observed reciprocal lattice points. We then find the optimal $A_{\boldsymbol{n}}$ and $B_{\boldsymbol{n}}$ that minimize the loss between the resultant $F'_{\boldsymbol{h}} = \frac{F_{\boldsymbol{h}}}{\sqrt{\epsilon_{\boldsymbol{h}}\Sigma''_{\boldsymbol{h}}}}$ and the standard Wilson distribution, leaving out 15% of the reflections for cross-validation to determine the optimal maximal value of components in $\boldsymbol{n}$.

Finally, we locally estimate $\Sigma_{\boldsymbol{h}}''' = \frac{\mathbb{E}(|\boldsymbol{F_h}|^2)}{\epsilon_{\boldsymbol{h}}}$ using $k$-nearest neighbor (KNN) estimation of $\mathbb{E}(|\boldsymbol{F_h}|^2)$ in reciprocal space, where $\epsilon_{\boldsymbol{h}}$ denotes the multiplicity as before. Starting from the structure factor amplitudes $F_{\boldsymbol{h}}'$ in the previous step, we compute $\Sigma_{\boldsymbol{h}}'''$ across 50-1,600 neighbors. We choose the optimal kernel size by cross-validation. We do a final scaling $E_{\boldsymbol{h}} = \frac{F_{\boldsymbol{h}}'}{\sqrt{\epsilon_{\boldsymbol{h}}\Sigma_{\boldsymbol{h}}'''}}$ to arrive at approximately normalized structure factors distributed according to the standard Wilson distribution (**Figure S11**).

#### Partial excitation

Perturbations are often applied to protein crystals to study protein dynamics. However, typically only a portion $p$ of the protein molecules in a crystal is driven from the ground state (GS) into the one or more excited states (ES). Here, we examine a model of such partial excitation introduced by Coppens et al.[15]. In this 'random distribution' model, the (complex) ON structure factors $\boldsymbol{F}^{on}$, i.e., the structure factors after perturbation is applied, depend on the ground-state structure factors $\boldsymbol{F}^{off} = \boldsymbol{F}^{gs}$ as well as the excited-state structure factors $\boldsymbol{F}^{es}$ so that

$$\boldsymbol{F}^{on} = p\boldsymbol{F}^{es} + (1-p)\boldsymbol{F}^{gs}$$

Under the assumptions described above (see **The bivariate Wilson distribution**), we observe that

$$\mathbb{E}(\boldsymbol{F}^{on}) = (1 - p + p \cdot r)\boldsymbol{F}^{gs} < \boldsymbol{F}^{gs},$$

since $p \in [0,1)$ and $r < 1$, such that, on average, ON amplitudes are slightly smaller than OFF amplitudes. To sensitively measure the effect of a perturbation on a crystal, often a difference map is constructed by subtracting related structure factor amplitudes, here $F^{on}$ and $F^{off}$, and combined with the OFF phases. As a result, naïve ON-OFF difference maps can be negatively correlated with the ground-state density. It is common to scale the related datasets to each other. With the multivariate Wilson model, we can analyze the effects of this choice on the difference map bias and ground-state correlation. In **Supplementary Notebook 6**, we compute two statistics to assess the quality of the scaled difference map $\boldsymbol{\Delta F}_{obs} = (kF^{on} - k'F^{off})\exp(i\varphi^{off})$, where $\varphi^{off}$ is the phase of the ground state: (1) the mean squared error between the scaled difference map and the true difference map $\boldsymbol{\Delta F}_{\mathbf{true}} = \boldsymbol{F}^{es} - \boldsymbol{F}^{gs}$ averaged over the unit cell, and (2) the covariance between the scaled difference map and the ground-state electron density. We find that placing the ON and OFF structure factors on the same scale (e.g., by choosing $\frac{k}{k'}$ as the median ratio of $F^{off}/F^{on}$ or $\sum_h F^{off} / \sum_h F^{on}$) effectively minimizes the covariance with the OFF density while keeping mean squared map error low and correlation with the true difference map near its maximum (**Figure S12**), justifying the practice of scaling related structure factor amplitudes to each other after merging.

#### Joint distributions of phases

Equation [5] also implies a statistical relationship between the phases of two data sets. Indeed, the difference of the calculated phases between refined *apo* and inhibitor-bound models follow the expected Von Mises distribution[6,16]

$$P(\Delta\varphi|E_1, E_2) = \frac{1}{2\pi I_0(z)} \exp(z\cos(\Delta\varphi))$$

with

$$z = \frac{2E_1E_2}{(1-r^2)}$$

We illustrate a fit of this model to the obtained phase differences in **Figure S13**.

### Supplementary Tables

| | NaI-soaked lysozyme at 24-ID-C |
|---|---|
| **PDB ID** | 9B7C |
| **number of passes** | 3 |
| **rotation (°)** | 720 |
| **Wavelength (Å)** | 1.03752 |
| **Resolution range (Å)** | 56.15 - 1.101 (1.14 - 1.101) |
| **Space group** | P 43 21 2 |
| **Unit cell** | 79.41 Å, 79.41 Å, 37.84 Å, 90°, 90°, 90° |
| **Total reflections** | 5,583,525 (44,497) |
| **Unique reflections** | 85,431 (1,781) |
| **Multiplicity** | 65.4 (13.1) |
| **Completeness (%)** | 91.24 (36.20) |
| **Mean I/sigma(I)** | 38.87 (1.44) |
| **Wilson B-factor (Å$^2$)** | 14.17 |
| **$R_{merge}$** | 0.05847 (1.882) |
| **$R_{meas}$** | 0.05887 (1.958) |
| **$R_{pim}$** | 0.00674 (0.523) |
| **$CC_{1/2}$** | 1.000 (0.480) |
| **CC*** | 1.000 (0.810) |
| **Mosaicity (°)** | 0.064±0.001 |
| **Figure of merit** | 0.40 |
| **Reflections used in refinement** | 45,167 (1,762) |
| **Reflections used for R-free** | 2,159 (81) |
| **R-work** | 0.1153 (0.2215) |
| **R-free** | 0.1267 (0.2231) |
| **$CC_{work}$** | 0.973 (0.859) |
| **$CC_{free}$** | 0.970 (0.889) |
| **$CC_{anom}$** | 0.655 (0.070) |
| **Number of non-hydrogen atoms** | 1,284 |
| **Macromolecules** | 1,176 |
| **Ligands** | 44 |
| **Solvent** | 86 |
| **Protein residues** | 129 |
| **RMS(bonds) (Å)** | 0.011 |
| **RMS(angles) (°)** | 1.05 |
| **Ramachandran favored (%)** | 99.21 |
| **Ramachandran allowed (%)** | 0.79 |
| **Ramachandran outliers (%)** | 0 |
| **Rotamer outliers (%)** | 0 |
| **Clashscore** | 2.1 |
| **Average B-factor (Å$^2$)** | 19.45 |
| **macromolecules** | 18.39 |
| **ligands** | 27.97 |
| **solvent** | 31.74 |

**Table S1.** Data collection, processing, and refinement statistics for the high-resolution monochromatic structure of NaI-soaked lysozyme. The refined model was used to phase the anomalous difference maps for Figure 3.

| Dataset | PanDDA peak height (σ) | multivariate Wilson scaling peak height (σ) |
|---|---|---|
| **P0115** | 35.29 | 37.41 |
| **P0116** | 8.41 | 8.72 |
| **P0123** | 14.45 | 7.67 |
| **P0124** | 20.72 | 16.64 |
| **P0131** | 9.61 | 16.06 |
| **P0132** | 19.73 | 15.92 |
| **P0137** | 19.02 | 16.65 |
| **P0138** | 28.94 | 24.72 |
| **P0139** | 13.79 | 14.24 |
| **P0142** | 6.96 | 6.32 |
| **P0148** | 11.18 | 13.16 |
| **P0159** | 37.16 | 38.41 |
| **P0161** | 41.49 | 45.87 |
| **P0163** | 19.21 | 14.48 |
| **P0178** | 26.14 | 48.31 |
| **P0179** | 20.88 | 21.74 |

**Table S2.** Comparison between PanDDA and multivariate Wilson analysis of the tallest ligand peak heights per dataset for the Mac1 fragment screening dataset.

### Supplementary Figures

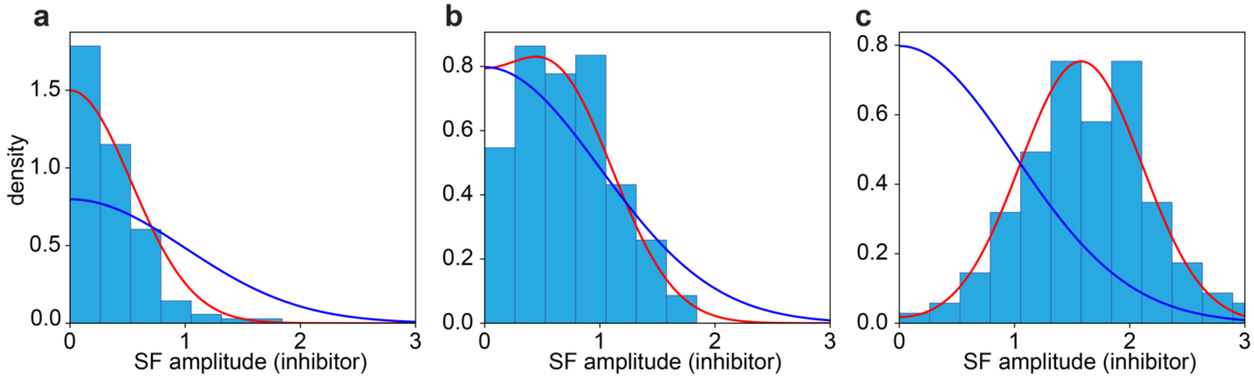


**Figure S1. Conditional distributions describe centric structure factor amplitudes.** Analogous to **Figure 1c-e**, histograms of *centric* structure factor amplitudes of PTP-1B in the presence of the TCS-401 inhibitor for reflections for which the structure factor amplitudes in the unliganded (apo) state fall within the **a)** 0-4$^{th}$, **b)** 48-52$^{th}$ and **c)** 92-96$^{th}$ percentile. Red: folded normal distributions parametrized by the mean apo structure factor amplitude per bin, and $r_{DW} = 0.85$. Blue: centric Wilson distribution.

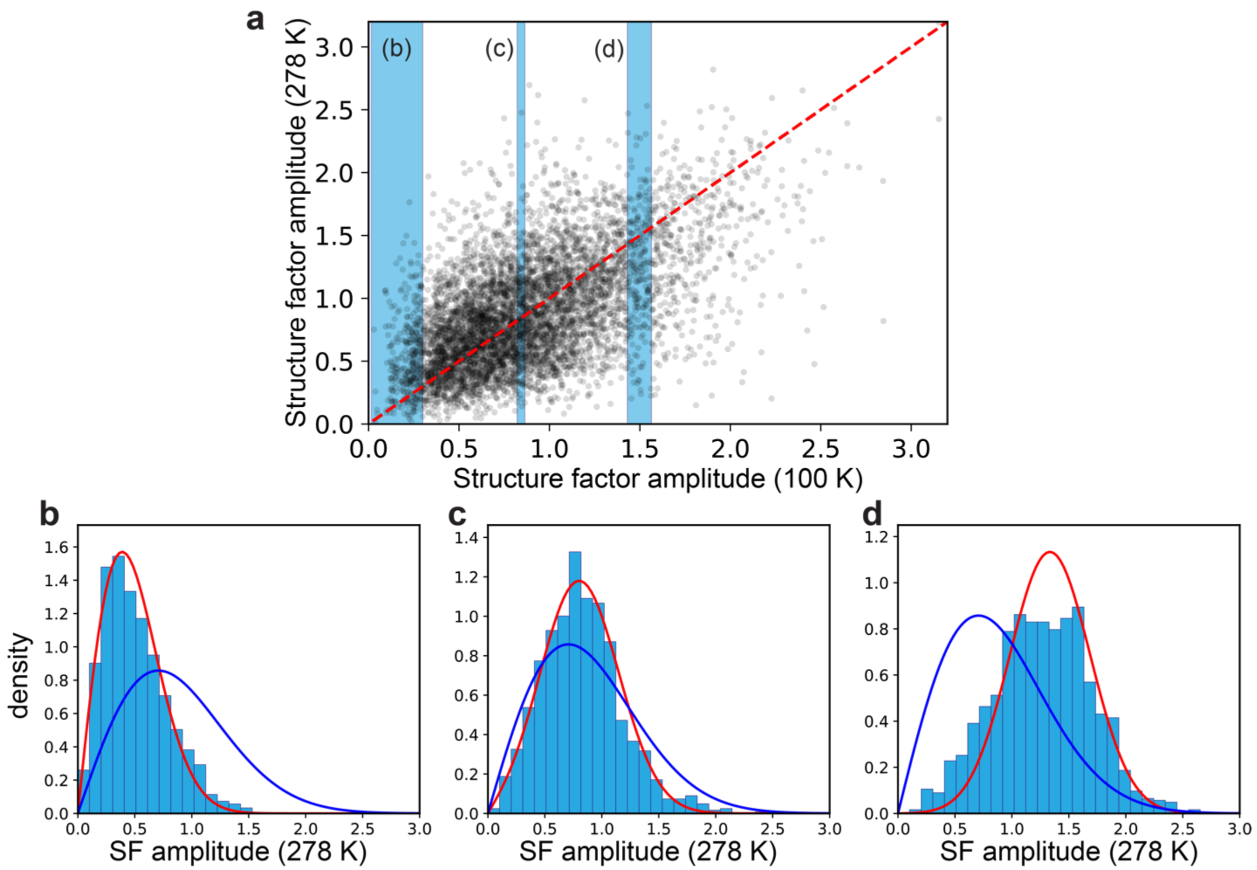


**Figure S2**. **The bivariate Wilson distribution for a pair of datasets across temperatures. a)** Scatter plot for a random subset of acentric structure factor amplitudes for two datasets of thaumatin, one measured at 100 K (PDB ID 5KVX[17]) and one measured at 278 K (PDB ID 5KW3[17]). Blue slices indicate data points for which histograms are shown in the next panels. **b-d)** Histograms for slices through shaded regions in panel **a** are better approximated by Rice distributions (red) parametrized by a double-Wilson $r$ (here, $r$ = 0.86) than by the Wilson distribution (blue).

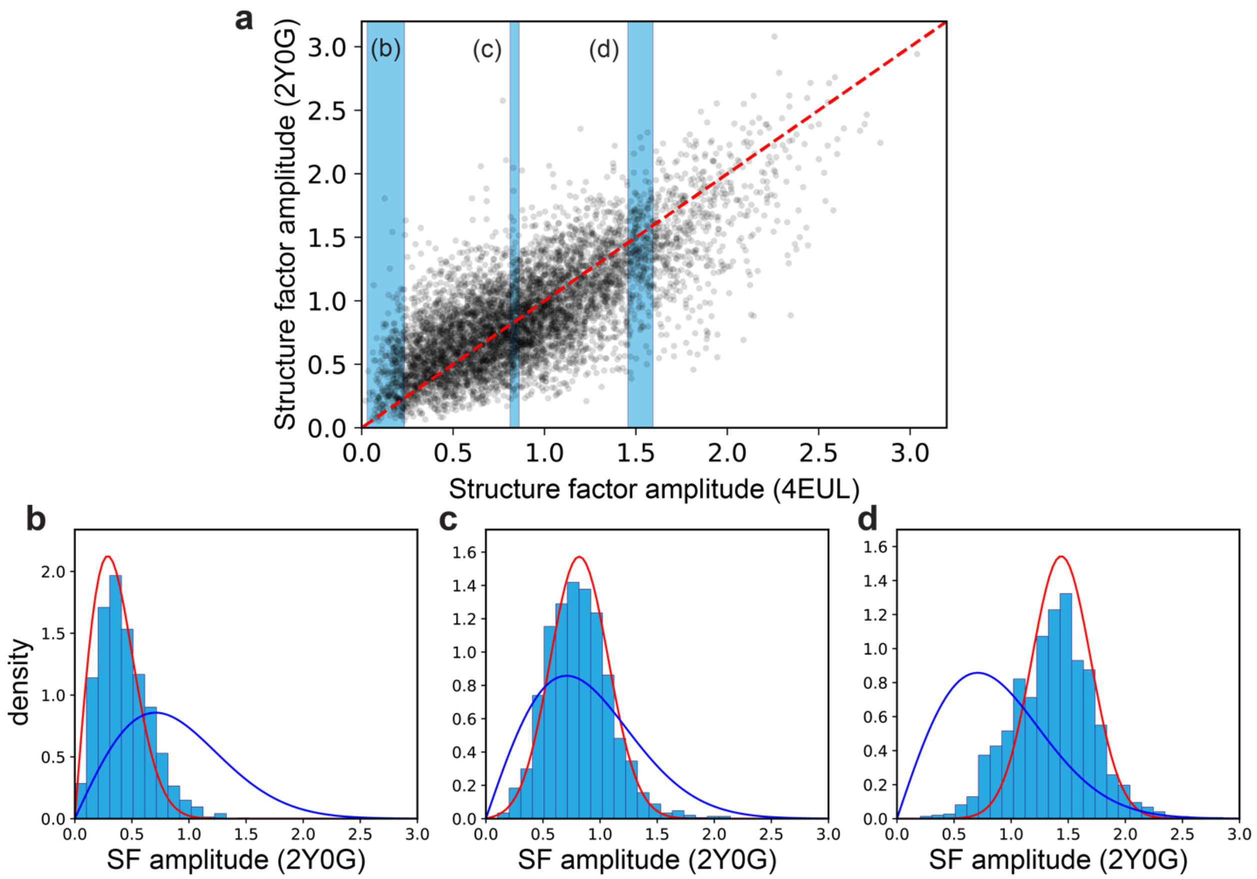


**Figure S3**. **The bivariate Wilson distribution for a pair of replicate datasets between labs. a)** Scatter plot for a random subset of acentric structure factor amplitudes for two datasets of eGFP, one reported in 2012 (PDB ID 4EUL[18]) and one reported in 2011 (PDB ID 2Y0G[19]). Blue slices indicate data points for which histograms are shown in the next panels. **b-d)** Histograms for slices through shaded regions in panel **a** are better approximated by Rice distributions (red) parametrized by a double-Wilson $r$ (here, $r = 0.93$) than by the Wilson distribution (blue).

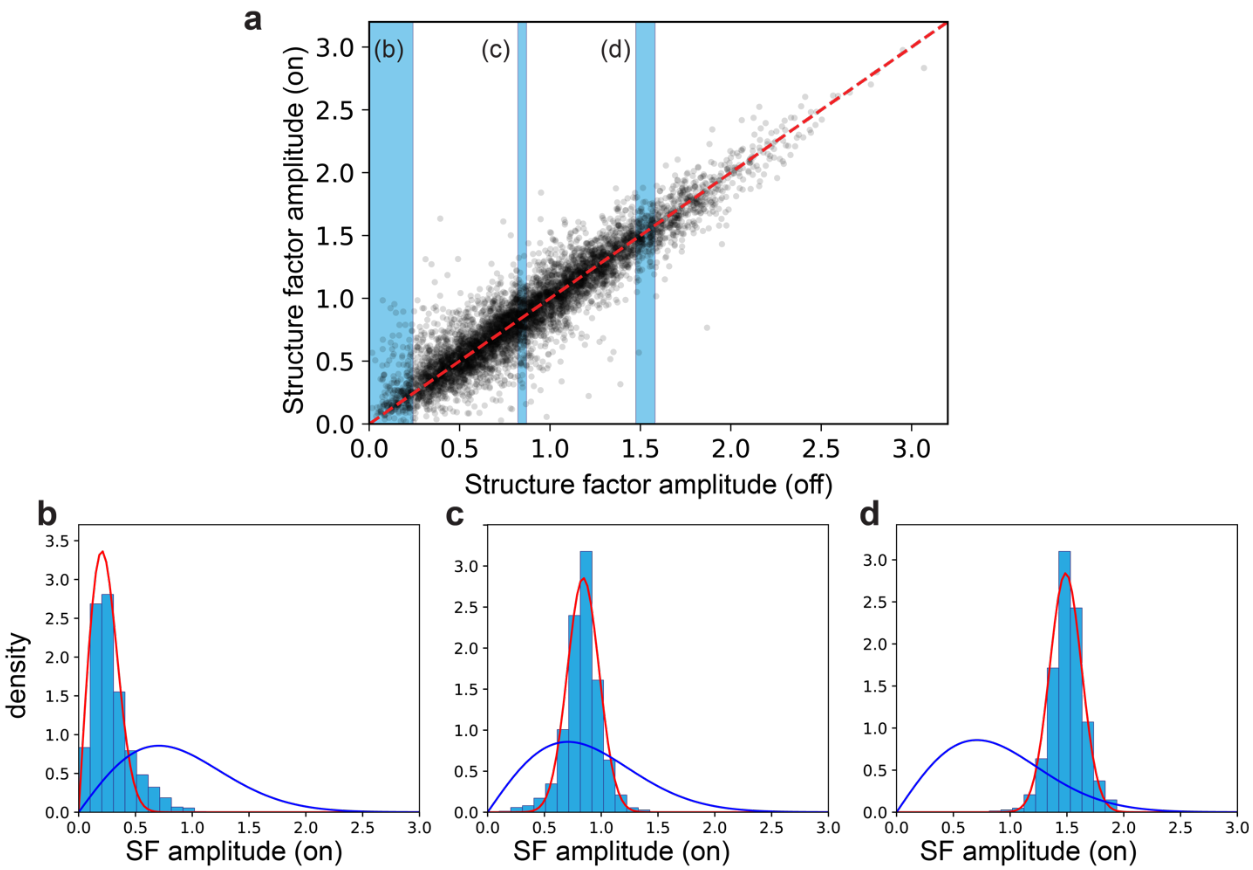


**Figure S4**. **The bivariate Wilson distribution for a pair of time-resolved datasets. a)** Scatter plot for a random subset of acentric structure factor amplitudes for two datasets of PYP, one measured without light illumination (PDB ID 1NWZ[20]) and one measured on a crystal cryotrapped after exposure to a 460 nm laser pulse (PDB ID 3PYP[21]). Blue slices indicate data points for which histograms are shown in the next panels. **b-d)** Histograms for slices through shaded regions in panel **a** are better approximated by Rice distributions (red) parametrized by a double-Wilson *r* (here, $r$ = 0.98) than by the Wilson distribution (blue).

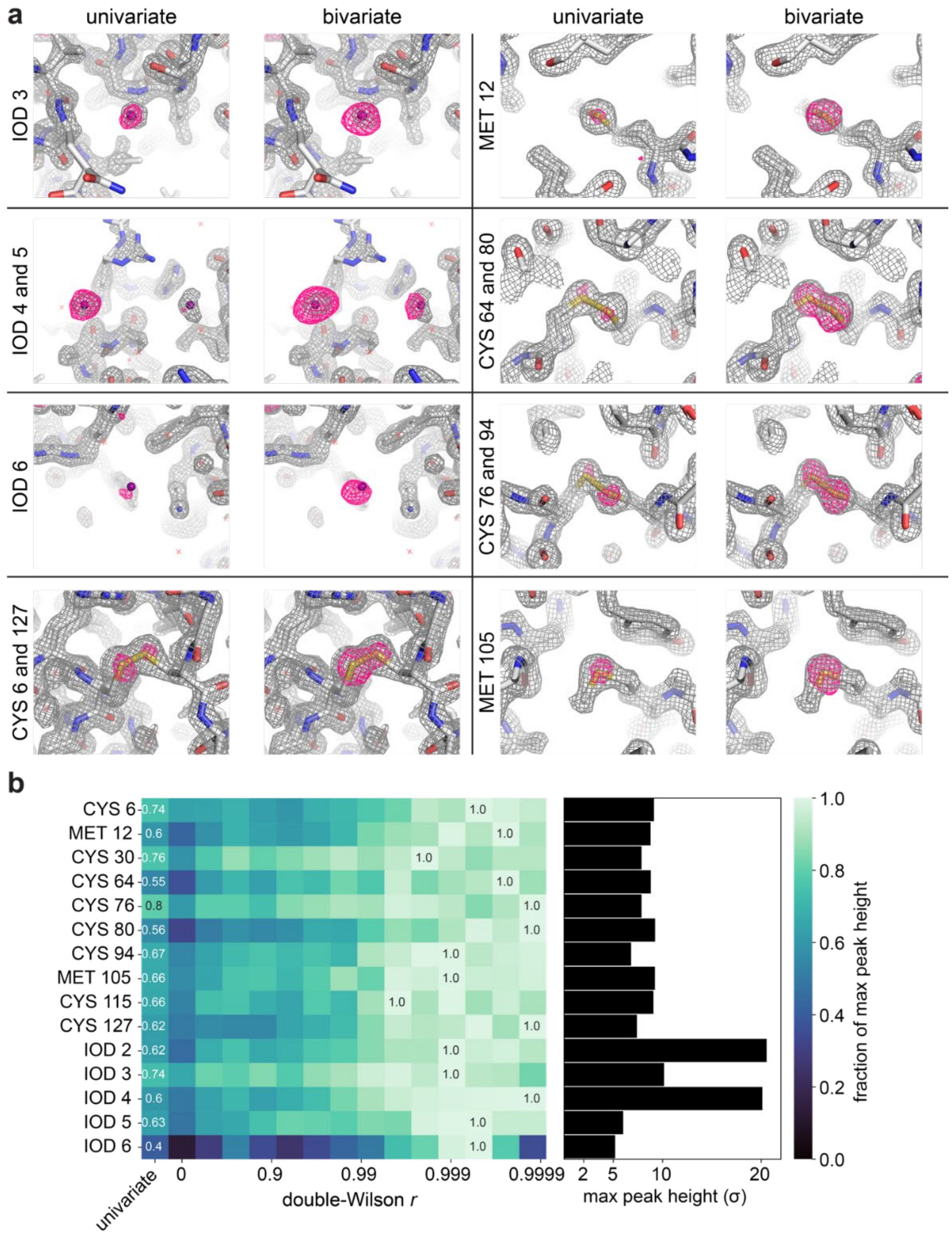


**Figure S5. NaI-soaked lysozyme anomalous omit peaks improve after scaling with a bivariate prior. a)** Comparisons between anomalous omit peaks merged with a bivariate prior and merged with a univariate prior. **b)** Peak heights of the anomalous difference peaks across *r*. **Left:** heatmap showing the fraction of the tallest peak across *r* for each anomalous scatterer in lysozyme. **Right:** absolute peak height of the tallest peak for each anomalous scatterer in lysozyme.

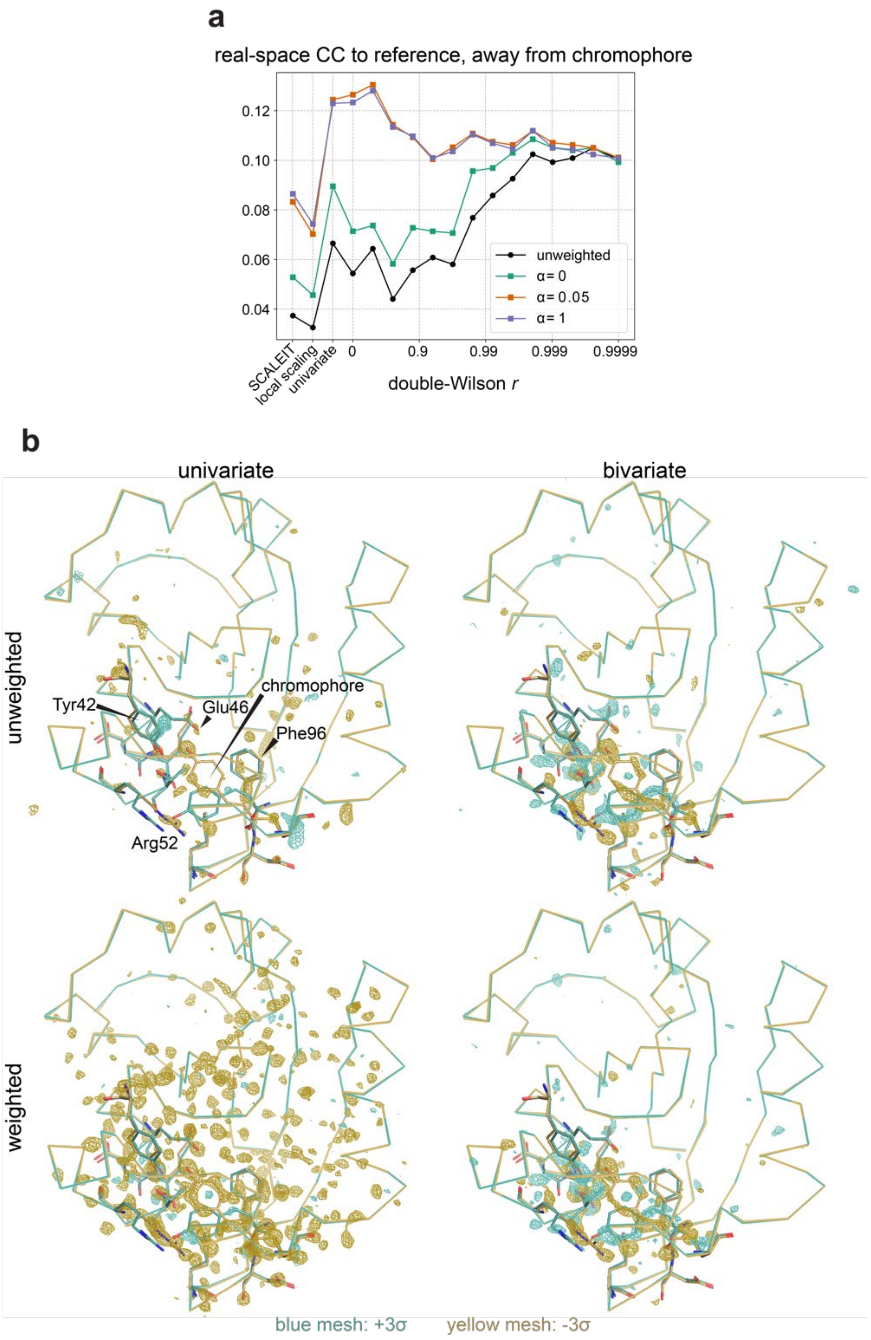


**Figure S6. PYP difference maps in regions far from the chromophore. a)** Real-space correlation coefficient between the observed and expected difference maps in the region more than 10 Å away from the chromophore. **b)** Weighted and unweighted difference maps scaled with a univariate and bivariate prior, *r*=0.99976. Difference maps are contoured to ±3σ and are colored as indicated. The teal and yellow models are the 2ms and off structures, respectively. The region within 10 Å of the chromophore is shown as sticks.

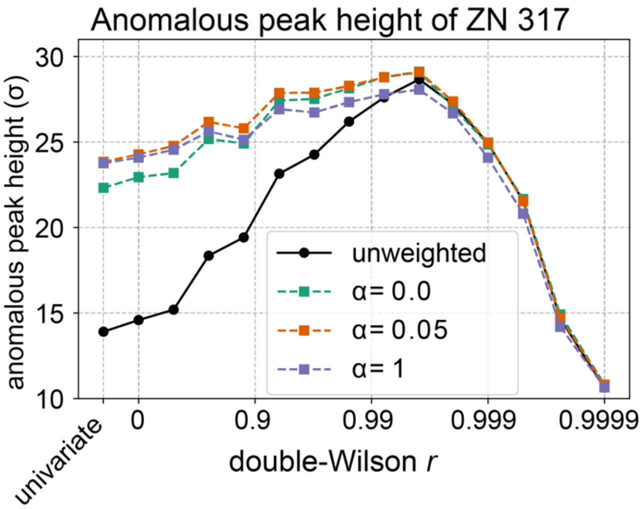


**Figure S7. Weighting anomalous differences in thermolysin omit maps. a)** The anomalous peak height of ZN 317 of thermolysin across *r*, for weighted (α=0, green; α=0.05, red; α=1, blue) and unweighted (black) anomalous differences.

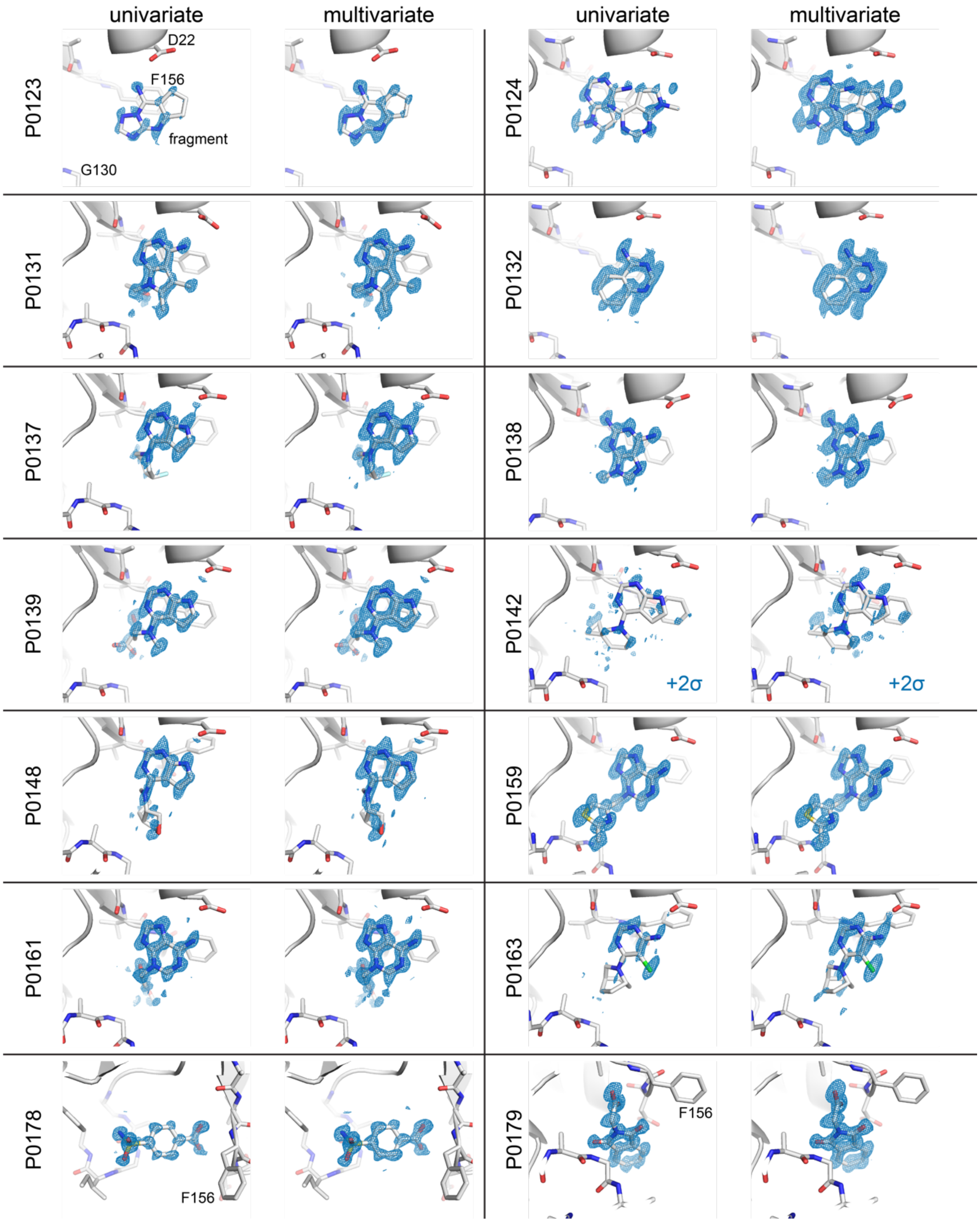


**Figure S8**. **Comparison of fragment screening difference maps scaled with a univariate and multivariate prior.** $F_{holo}$ - $F_{apo}$ difference maps from fragment screening of Mac1, scaled with a univariate and multivariate prior. All maps carved to 1.5 Å of the ligand and contoured to +3σ unless indicated.

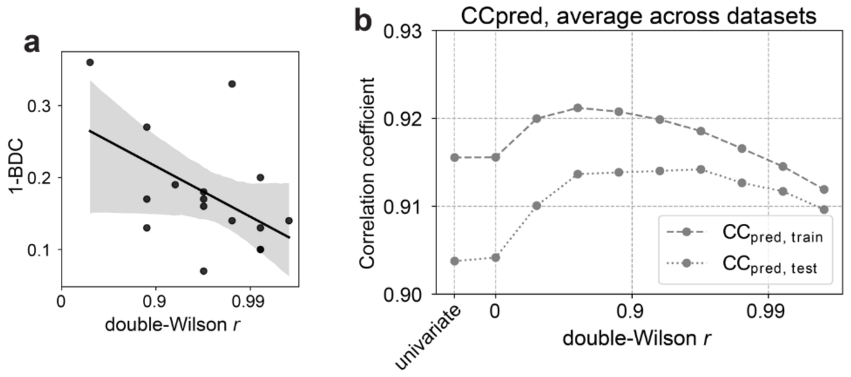


**Figure S9. Fragment screening dataset statistics. a)** plot of the optimal double-Wilson *r* value for each dataset, against the PanDDA 1 minus background data correction parameter, a proxy for the combined effects of occupancy and crystal idiosyncracies (see **Results**). The x-axis is on a log scale and the Pearson $r = 0.505$, with $p < 0.05$. **b)** Average $CC_{pred}$ across the *holo* and *apo* datasets for each value of the double-Wilson *r*.

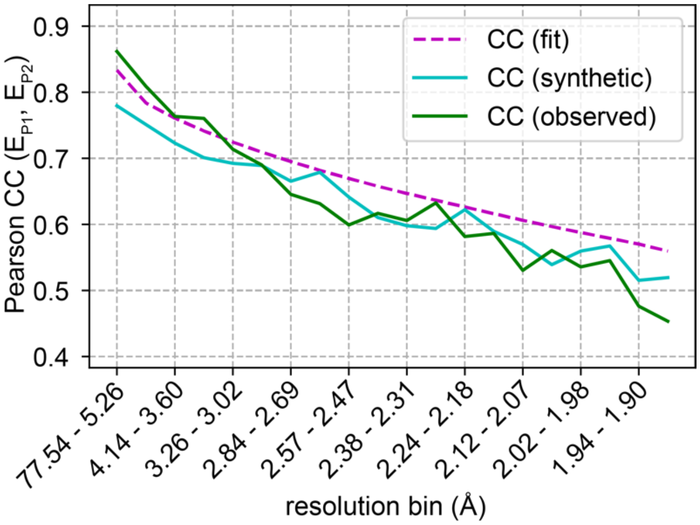


**Figure S10. Resolution dependence of the correlation coefficient (CC) between related datasets.** Green line: resolution-binned correlation coefficients between normalized structure factor amplitudes of PTP-1B apo and bound to the TCS-401 inhibitor. Magenta line: fit CC, $a = 0.91$ and $b = 0.71$ describing the inferred correlation between the true structure factor amplitudes (**Supplementary Information, "The bivariate Wilson distribution").** Cyan line: CC between two synthetic datasets with correlations described by the same parameters, combined with structure factor amplitudes and measurement errors drawn from their observed values.

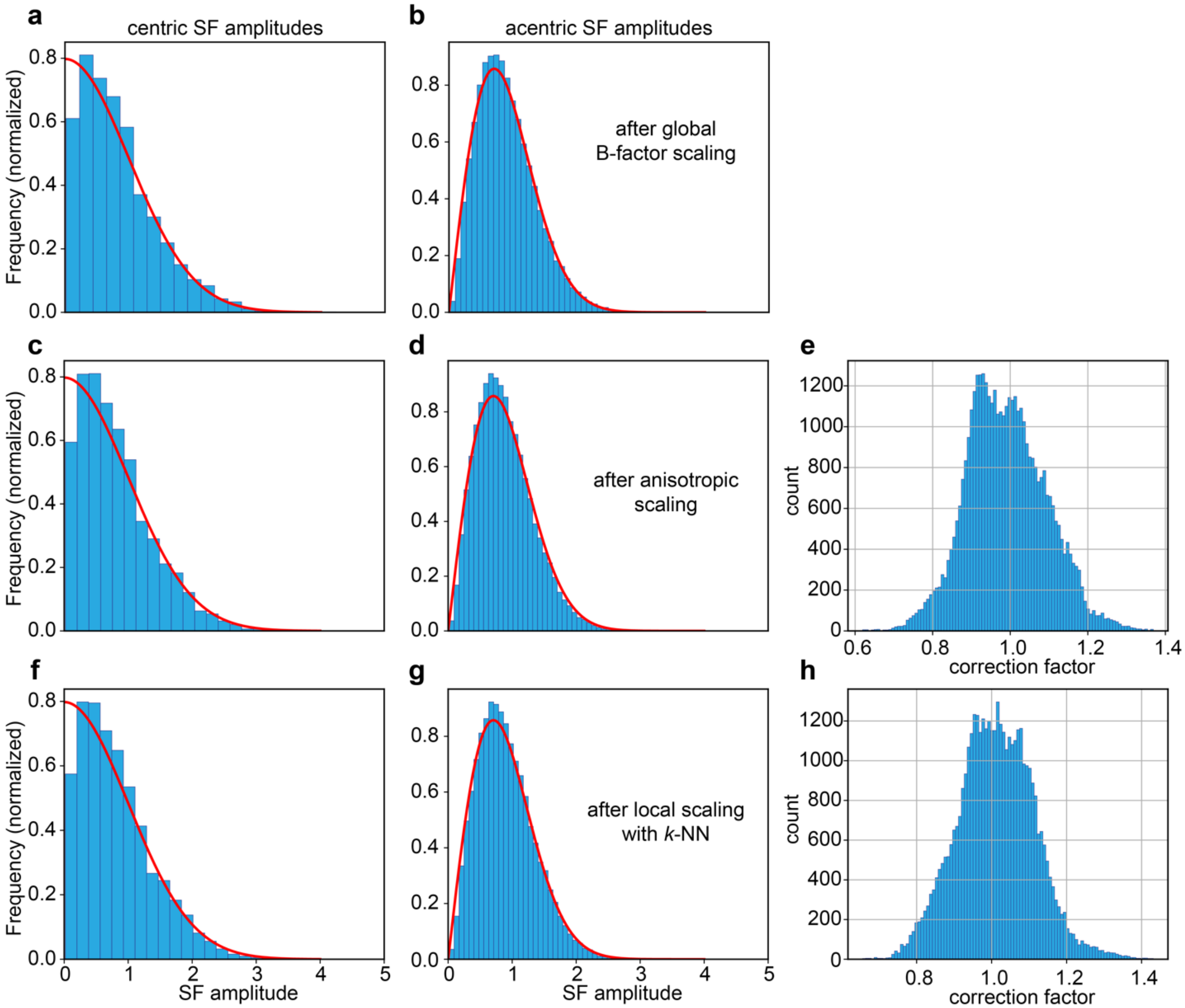


**Figure S11. Normalization and Wilson statistics of structure factors. a-b)** Distribution of structure factor amplitudes after global B-factor scaling for centric (**a**) and acentric (**b**) structure factor amplitudes. Red curves: expected distribution of normalized structure factor amplitudes under the Wilson distribution. **c-d)** Distribution of structure factor amplitudes after simple anisotropic scaling for centric (**c**) and acentric (**d**) structure factor amplitudes. Red curves again represent the Wilson distribution. **e)** Distribution of correction factors for a Fourier series correction of structure factor amplitudes (see **Supplementary Information**). **f-g)** Distribution of normalized structure factor amplitudes after additional correction based on *k*-nearest neighbor ridge regression for centric (**f**) and acentric (**g**) reflections. Red curves again represent the Wilson distribution. Data for apo PTP-1B[22]. **h).** Distribution of correction factors relative to naïve anisotropic scaling.

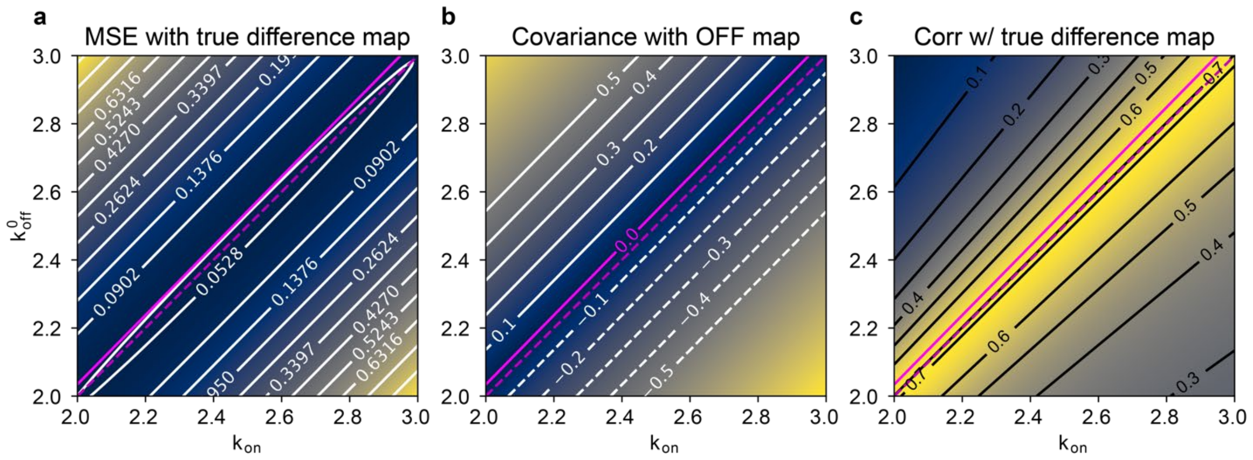


**Figure S12. Negative correlation with the unperturbed state electron density can be removed without degrading difference map quality. a)** the mean squared error (MSE) between the scaled difference map $(kF^{on} - k'F^{off})\exp(i\phi^{off})$ and the true difference map between ground state and excited state $(\boldsymbol{F^{es}} - \boldsymbol{F^{gs}})$ plotted across $k$ and $k'$.**b**) The covariance between the scaled difference map and the unperturbed (OFF) electron density map**. c**) The Pearson correlation between the scaled difference map and the true difference map. Magenta solid lines: the line of 0 covariance between the scaled difference map and the unperturbed map $\boldsymbol{F^{off}}$. Magenta dashed lines: $k = k'$. See **Supporting Information** for notation. White lines: isocontour lines for each plot with contour isovalues shown (solid contour lines for positive values, dashed contour lines for negative values).

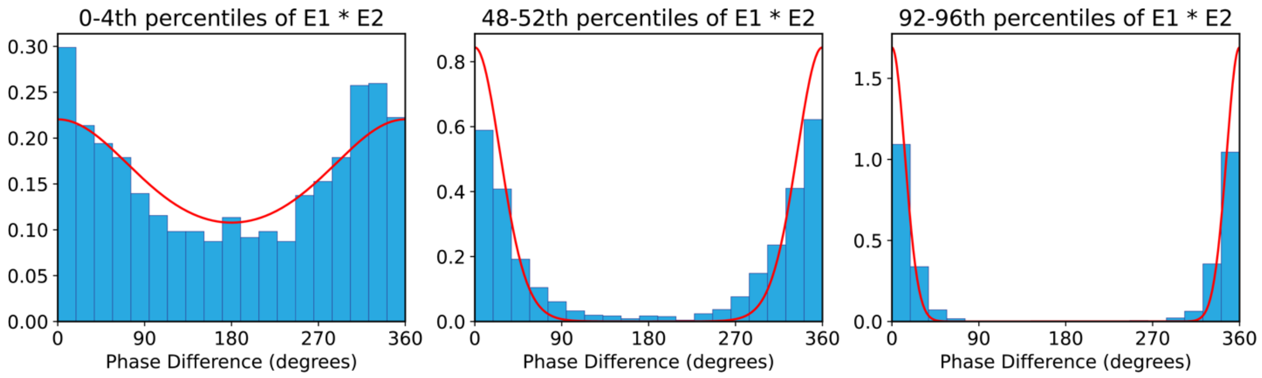


**Figure S13. Von Mises statistics of phase differences.** Phase differences for acentric reflections (blue bars) calculated from structure factors of PTP-1B in the absence and presence of the TCS-401 inhibitor. Von Mises distribution (red line) fit to the phase differences.
